# Incorporating discovery and replication GWAS into summary data Mendelian randomization studies: A review of current methods and a simple, general and powerful alternative

**DOI:** 10.1101/2023.01.12.523708

**Authors:** Ninon Mounier, David S. Robertson, Zoltán Kutalik, Frank Dudbridge, Jack Bowden

## Abstract

Mendelian Randomization (MR) is a popular method for using genetics to estimate the causal effect of a modifiable exposure on a health outcome. Single Nucleotide Polymorphisms (SNPs) are typically selected for inclusion if they pass a genome-wide significance threshold in order to guarantee that they are strong genetic instruments, but this also induces Winner’s curse, as SNP-exposure associations tend to be overestimated. In this paper, we consider how to combine SNP-exposure data from discovery and replication samples using two-sample and three-sample approaches to best account for Winner’s curse, weak instrument bias, and pleiotropy within a summary data MR framework, using only GWAS summary statistics. After reviewing several existing methods, that often correct for Winner’s curse at the individual SNP level, we propose a simple alternative based on the technique of regression calibration that enacts a global correction to the causal effect estimate directly. This approach does not only correct for Winner’s curse, but also simultaneously accounts for weak instruments bias. Regression calibration can be used with a wide range of existing MR methods, including pleiotropy-robust methods such as median-based and mode-based estimators. Extensive simulations and real data examples are used to illustrate the utility of the new approach. Software is provided for users to implement the method in practice.

**Author Summary:** Mendelian randomization is a method to explore causation in health research which exploits the random inheritance of genes from parents to offspring as a ‘natural experiment’. It attempts to quantify the effect of intervening and modifying a health exposure, such as a person’s body mass, on a downstream outcome such as blood pressure. Causal estimates obtained using this method can be strongly influenced by the set of genes used, or more specifically, the rationale used to select them. For example, selecting only genes that are strongly associated with the health exposure can induce bias due to the ‘Winner’s curse’. Unfortunately, using genes with a small association can lead to so-called ‘weak instrument’ bias leading to a no-win paradox. In this paper, we present a novel approach based on the technique of regression calibration to de-bias causal estimates in an MR study. Our approach relies on the use of two independent samples for the exposure (discovery and replication) to estimate the amount of bias that is expected for a specific set of genes, so that causal estimates can be re-calibrated accordingly. We use extensive simulations and applied examples to compare our approach to current methods and provide software for researchers to implement our approach in future studies.

## 1 Introduction

Mendelian randomization (MR) [1] is an increasingly popular technique for exploiting genetic data to estimate the causal effect of a modifiable exposure on a health outcome. The appeal of MR is that it can in principle provide consistent causal estimates without the need to adjust for all variables which confound the exposure-outcome relationship, provided that the genetic variants used satisfy the Instrumental Variable (IV) assumptions. These are encoded within the causal diagram in Figure 1. Here, *G*_*j*_ represents a genetic variant - usually a Single Nucleotide Polymorphism (SNP); *X* represents a health exposure; *Y* represents the health outcome and *U* represents unmeasured confounders of *X* and *Y*.

**Figure 1:**
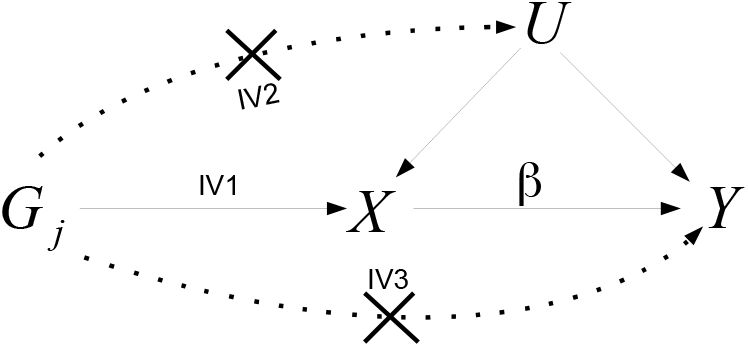
Causal diagram representing the assumed relationship between a genetic variant G_j_, exposure X and outcome Y within a Mendelian randomization study.

The *G*_*j*_ → *X* arrow denotes that *G*_*j*_ predicts *X*. This can be empirically verified from the data and is referred to as the ‘relevance’ assumption (IV1). The lack of an arrow from *G*_*j*_ to *U* (IV2) or from *G*_*j*_ to *Y* directly (IV3) denotes that *G*_*j*_ only affects *Y* through *X*. This is referred to as the ‘exclusion restriction’ assumption. If these assumptions are satisfied, a non-zero association between *G*_*j*_ and *Y* can be used to reject the null hypothesis of no causal effect of *X* on *Y*. Further linearity and homogeneity assumptions are required in order to estimate the causal effect, *β*, of a unit increase in *X* on *Y*, which is denoted by the arrow *X* → *Y* in Figure (1).

One reason for a surge in popularity of MR methods is the breadth and scope of designs now available. In a traditional ‘one-sample’ analysis, individual-level data from a single cohort is used to construct a genetically predicted exposure, 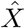. When regressed on the outcome *Y*, the coefficient for 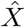 is then taken as a causal effect estimate. An alternative approach is ‘two-sample’ summary data MR, in which summary SNP-exposure and SNP-outcome associations from two genome-wide association studies (GWAS) are combined together using the principles of meta-analysis to provide a causal estimate [2, 3]. With the number of available GWAS summary statistics continuously increasing, coupled with easy-to-use data and analysis pipelines such as MR-base [4], two-sample MR has steadily gained popularity in the past decade. Figure 2-A provides a simple overview of the two-sample summary data approach and the important assumptions upon which it rests.

The SNP statistics that are needed for the estimation of the causal effect are described in Figure 2-B. They can be affected by different sources of bias, which could lead to biased inference on the existence and magnitude of the true causal effect. A major focus of research in MR has been the development of methods to mitigate the consequences of SNPs that violate the exclusion restriction by exerting a pleiotropic effect on the outcome not through the exposure [5, 6, 7, 8]. Some methods allow for a weaker version of the exclusion restriction - the Instrument Strength Independent of Direct [pleiotropic] Effect (InSIDE) assumption - to be violated and can still in theory deliver unbiased causal estimates [9]. However, InSIDE is generally violated when the magnitudes of the pleiotropic effects are correlated with instrument strengths due to acting via a heritable confounder. This can lead to causal estimates that are biased towards the observational association between *X* and *Y*. Pleiotropy-robust estimators, such as median-based estimators that rely on the ‘majority’ rule (at least 50% of the instruments are valid [8]) or mode-based estimators that rely on the ‘plurality’ rule (non-pleiotropic instruments are the most frequent [6]) can be reliably used when the InSIDE assumption is violated.

**Figure 2:**
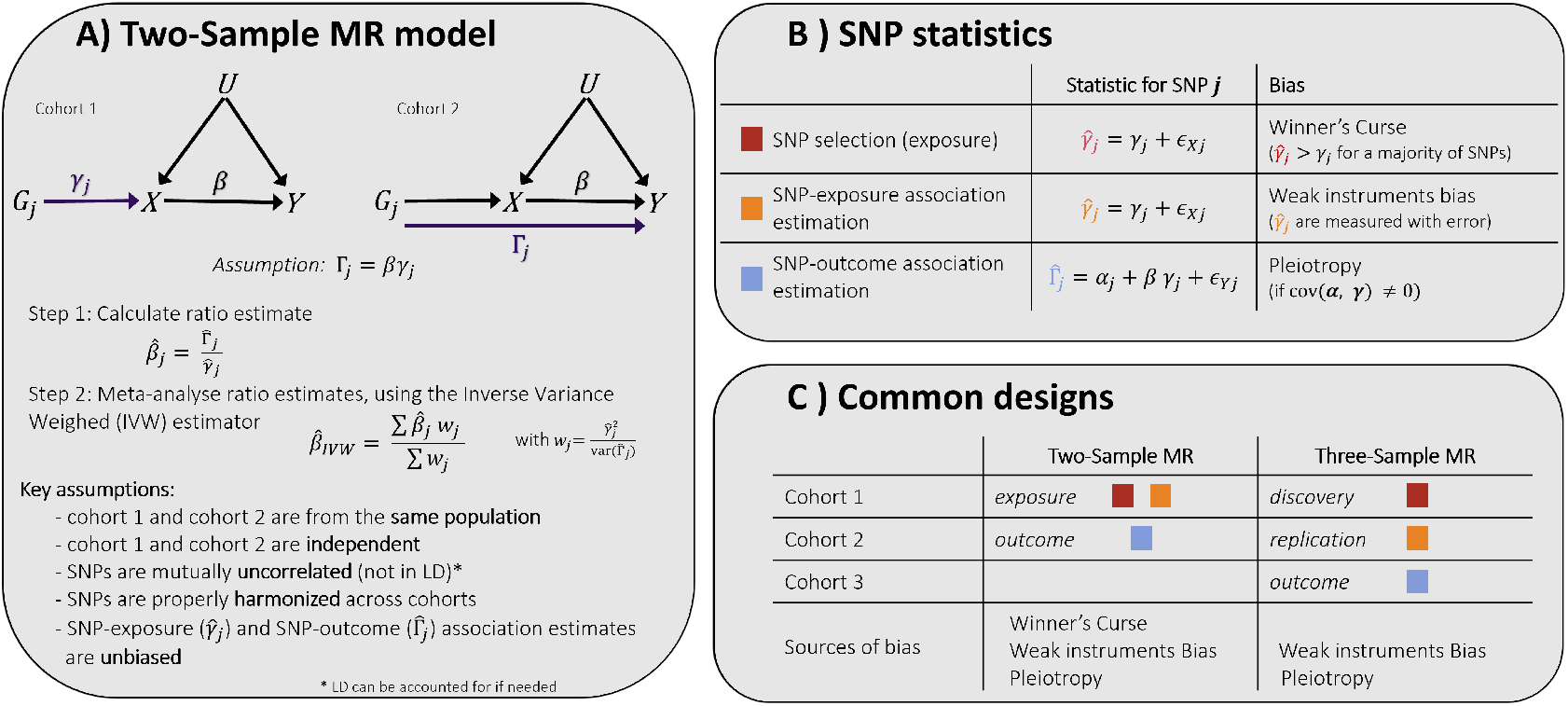
Panel A describes the two-sample MR model and the key assumptions it relies on. Panel B summarizes the different statistics needed to perform the analysis and the biases that might affect them. Panel C compares the two-sample and the three-sample designs.

Most MR methods make the assumption that SNP-exposure association estimates are so precise that their uncertainty is negligible, or that the ‘NO Measurement Error’ (NOME) assumption is satisfied [8]. NOME can, to all intents and purposes, be forced to hold by selecting SNPs as instruments based on a strict p-value criterion, such as the ubiquitous genome-wide significance threshold of *p* ≤ 5*×*10^−8^. If this selection phase is not done, or no SNP meets this threshold in the data at hand, then MR estimates are typically affected by weak-instrument bias, which is equivalent to regression-dilution bias towards the null in a two-sample analysis. Some methods such as MR-RAPS [10] and Radial MR [11] incorporate an internal weak instrument bias correction to mitigate this effect.

One consequence of selecting instruments based on statistical significance, and then using their estimated associations in the MR analysis, is Winner’s curse [12]. This refers to the phenomenon that SNP-exposure associations tend to be overestimated when a threshold is used to select instruments. The issue was first discussed in detail in the two sample summary data MR context by Zhao et al. [10]. Their solution was to use two sets of GWAS summary statistics for the exposure: one to select instruments using discovery data, and the other to re-estimate or ‘replicate’ the SNP-exposure association estimates to be used in the actual MR analysis. This has become known as the ‘three sample’ design strategy (Figure 2-C) for obvious reasons. It typically leads to the inclusion of weaker genetic instruments, hence the specific need for weak instrument bias adjustment. More sophisticated approaches to de-bias the discovery SNP-exposure association estimates have also been proposed in the general GWAS literature [12, 13, 14, 15, 16], but are not frequently used because the focus is not on estimation but rather on detecting true non-zero associations and associated issues of error rate control.

In this paper, we will consider different strategies to combine discovery and replication data to properly account for weak instrument bias and Winner’s curse. We will also introduce a novel approach based on the general principle of regression calibration [17, 18]. This approach is naturally robust to both weak instrument bias and Winner’s curse, uses an intuitive correction to re-calibrate the causal effect estimate, and can be applied to various MR methods, including pleiotropy-robust ones. We compare the performance of the different strategies using both simulated data and real data from UK Biobank (UKB) [19] and provide easy-to-use software to implement all of the methods in practice.

## 2 Methods

We start by assuming the following data-generating model for the summary genetic associations used in the MR analysis:

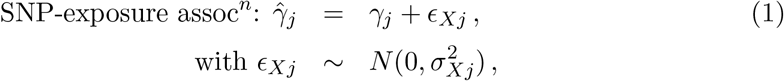

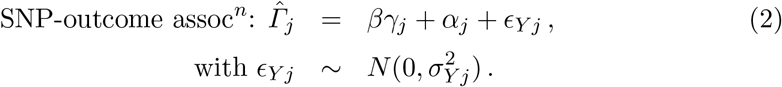

Here, 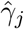 represents the set of standardised SNP-exposure association estimates, with variance 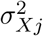 (directly related to the exposure sample size, 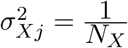) and 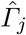 represents the set of standardised SNP-outcome association estimates, with variance 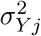 (directly related to the outcome sample size, 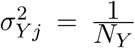). The value of 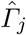 is influenced by the true SNP-exposure association (*γ*_*j*_), the causal effect (*β*), and the (direct) pleiotropic effect of SNP *j* on *Y* not through the exposure (*α*_*j*_). We will initially assume that the pleiotropy is ‘balanced’. That is, the mean pleiotropic effect is zero and independent of the SNP-exposure association, so that the sample covariance of the underlying pleiotropic effect and instrument strength parameters is:

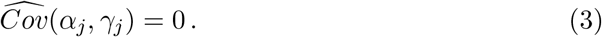

In this case, *α*_*j*_ can be treated as a zero centred random effect in model (2). Model 2 also tacitly assumes that the causal effect is constant across levels of the exposure predicted by the instruments. Although this has been relaxed in extended models [20, 21], we will not consider this here.

If we assume that there is correlated (and hence unbalanced) pleiotropy, for example in the case of the existence of a heritable confounder *U* that affects both X (with an effect *q*_*X*_) and Y (with an effect *q*_*Y*_), the data generating model can be re-written as follows:

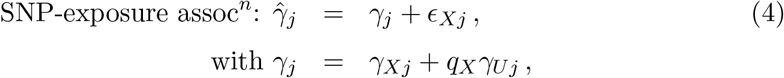

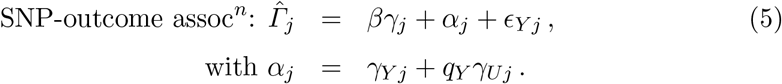

Here *γ*_*Xj*_ represents the direct effect of the SNP on the exposure, while *γ*_*Uj*_ corresponds to its effect on the confounder. *γ*_*Y j*_ represents the direct effect of the SNP on the outcome. In this case, if both *q*_*X*_ and *q*_*Y*_ are non-zero, the pleiotropic effect of SNP *j* on Y is not independent of the SNP-exposure association and we have

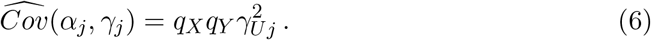

The ratio estimate for the causal parameter *β* using SNP *j* only is

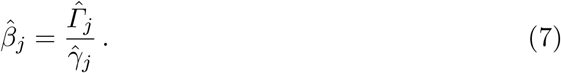

In the following sections, we will assume that SNP-exposure association estimates are coming from two independent samples: a discovery sample (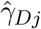, with sample size *N*_*D*_ and variance 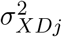), used for instrument selection, and a replication sample (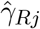, with sample size *N*_*R*_ and variance 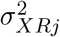), used for the estimation of the causal effect.

Standard two-sample MR analyses are susceptible to bias due to Winner’s curse and weak instruments. Both result in a bias towards the null for different reasons: Winner’s curse leads to over-estimation of the *j*th Wald ratio denominator in (7) and weak instruments induce regression-dilution because across all SNPs, 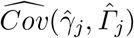 is less than 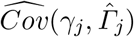.

Instrument strength is nicely measured using the mean *F* statistic, 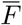, across the entire set of *L* SNPs,

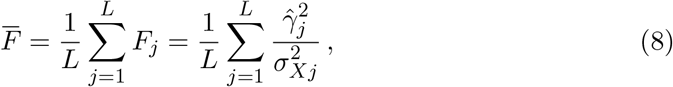

since the resulting weak-instrument bias dilution in MR estimate 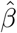 is equal to 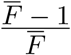 [10]. When only genome-wide significant SNPs (*p*-value ≤ 5*×*10^−8^, *T* ≈ 5.45) are used each SNP’s individual F-statistic, *F*_*j*_, is greater than approximately 30. Although this superficially alleviates weak instrument bias it does so at the cost of inducing Winner’s curse.

In the remainder of this section we briefly describe existing approaches to combine discovery and replication data in an effort to mitigate the effects of Winner’s curse and/or weak instrument bias (Table S1). Then we provide further details on our novel regression calibration approach. Finally, we present the simulation designs and describe the data used for applied examples.

### 2.1 Existing approaches

#### 2.1.1 Naive IVW estimate

The most common approach to combining causal estimates across SNPs is to perform an inverse variance-weighted (IVW) meta-analysis:

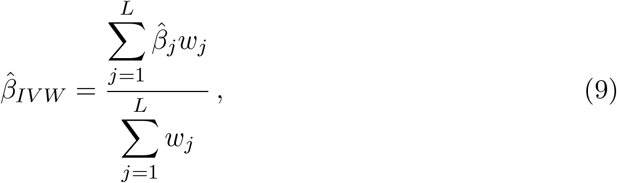

where *L* is the number of instruments used and 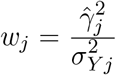 represents the inverse variance of 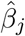, under the simplifying assumption that the uncertainty in the SNP-exposure association estimate is negligible (the NOME assumption). This naive implementation of the IVW approach does not separate out discovery and replication data; both are meta-analysed in order to work with a single combined exposure sample, using the largest sample size available. We denote the combined SNP-exposure association estimate as 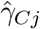, with sample size *N*_*C*_ = *N*_*D*_ + *N*_*R*_ and variance 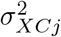. As previously discussed, using combined data increases the precision of the association estimates. This reduces but does not eliminate weak instrument bias and Winner’s curse. We use the IVW implementation from the `twoSampleMR` R-Package [22].

#### 2.1.2 IVW using a three-sample design

The simplest way to avoid Winner’s curse is to use the three-sample design proposed by Zhao et al. [10]. This requires access to a discovery and a replication sample. Instruments are selected based on their association in the discovery sample (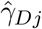 and 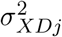) but only the SNP-exposure associations from the replication sample 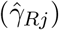 are used to estimate their ratio estimate in equation (7). The ratio estimates can then be combined using IVW (9), and again we use the implementation from the `twoSampleMR` R-Package to do so [22]. While this approach removes Winner’s curse, causal effect estimates are biased by weak instruments, since replicated SNP-exposure associations will generally have far lower *F* statistics.

#### 2.1.3 MR-RAPS using three-sample design

As illustrated in the previous section, when undertaking a three-sample design, causal effect estimates are still susceptible to weak instrument bias. To account for weak instruments, we use the MR-RAPS method [10] with the unbiased replication estimates (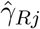 and 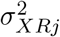) for instruments selected using discovery data (three-sample design). We used the robust, over-dispersed implementation from the `mr.raps` R-Package [10], incorporating the Huber loss function. The causal effect estimate obtained from this approach accounts for both Winner’s curse and weak instrument bias. Although its use of a robust loss function can mitigate the effects of highly pleiotropic SNPs by penalizing their weight in the analysis, it still heavily relies on the assumption of balanced pleiotropy (and hence InSIDE).

#### 2.1.4 MR-RAPS using a conditional likelihood

MR-RAPS can also be used in combination with other methods to account for Winner’s curse. It is for example possible to use a conditional likelihood approach to de-bias the SNP-exposure associations (see Supplementary A. for details), and this has been suggested multiple times in the literature, see for example Zhong and Prentice [12] and Xiao and Boehnke [13] and more recently Palmer and Pe’er [14]. We will refer to this as the ZP approach from now on. As this does not require separate discovery and replication samples, we can again use the combined data. The SNP-exposure association 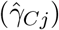 can then be corrected (using the implementation of the `gwas-winners-curse` R-Package [14]) to obtain a bias-reduced estimate of the effect of the SNP on the exposure, 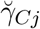. MR-RAPS can then be used with the corrected SNP-exposure association estimates (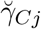 and the corresponding variance 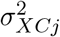) for instruments selected using the combined data, to deal with both Winner’s curse and weak instrument bias, with the aforementioned limitations.

#### 2.1.5 MR-RAPS using UMVCUE

An alternative method combining the discovery and replication data to obtain an efficient, unbiased estimate of the SNP-exposure associations is to employ Rao Blackwellisation, a technique popularised in the adaptive trial literature by Bowden and Glimm [23] and subsequently applied to the GWAS setting by Bowden, Dudbridge and Robertson [15, 16]. This method calculates the expected value of the unbiased replication estimate 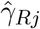, conditional on a complete sufficient statistic for *γ*_*j*_ and conditional on selection (for a selection threshold *T*). The resulting quantity is known as the Uniformly Minimum Variance Conditionally Unbiased Estimator (UMVCUE), denoted by 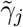. Details about how to obtain the UMVCUE estimator and its corresponding variance 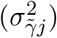 in MR analyses are available in Supplementary B.

It is then possible to use MR-RAPS with the UMVCUE SNP-exposure association estimates 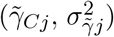 for instruments selected using discovery data only. This two-step correction accounts for both Winner’s curse and weak instrument bias.

#### 2.1.6 MRlap

The MRlap approach models the entire instrument selection process whilst accounting for exposure effect estimation due to weak instruments and Winner’s curse, as well as accounting for sample overlap if present (although this final feature is not relevant to the scope of this paper) [24]. To do so, it assumes a spike-and-slab distribution of the genetic architecture of the exposure, and relies on different techniques such as LD-Score regression to estimate the relevant parameters. It can be used with combined data as it does not need discovery and replication samples, but it does require genome-wide information about SNP-exposure and SNP-outcome associations (not only instruments) to estimate the underlying parameters needed for the correction.

### 2.2 Regression calibration

We now consider an adaption that can be made to the naive IVW estimate, that can simultaneously remove weak instrument bias and Winner’s curse. We first note that the bias of each SNP-exposure association is not of interest *per-se*. We only care about its effect on the overall IVW estimate in (9).

One way of estimating the overall bias for a given set of instruments is to use them to estimate the causal effect of the exposure on itself, using discovery and replication data:

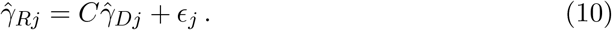

In other words, we can use the discovery sample as the exposure (to select instruments and estimate the SNP-exposure associations) and the replication sample as the outcome, in a standard two-sample MR IVW framework, to obtain the best fitting (or least-square) slope *Ĉ*:

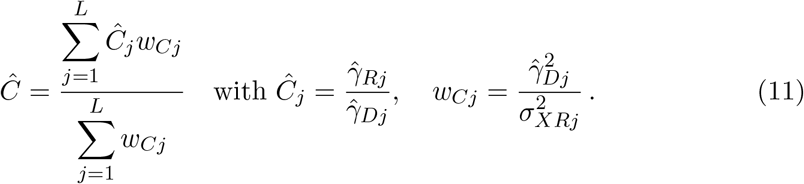

The slope *C* corresponds to the causal effect of a trait on itself and is expected to be 1 in the absence of bias. We propose to use the slope estimate to correct the naive IVW estimate in (9), by scaling it up by a factor of 1*/Ĉ*. That is, we calculate

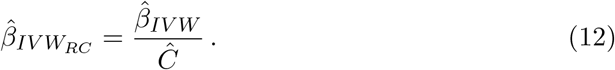

This only works if 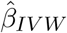 is obtained using only the discovery sample (to select instruments and estimate SNP-exposure associations) so that the same set of instruments is used in (9) and in (11). This re-calibration of the naive IVW causal effect estimate does not explicitly model Winner’s curse nor weak instrument bias, it estimates from the data the amount of dilution that is induced by the use of the specific set of instruments derived from the discovery sample. The explanatory variable in both regressions is 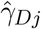 and the bias then cancels out. In this case, the replication sample is only used to obtain *Ĉ*. The variance of the estimate 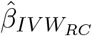 is approximated using the first term of a Taylor series expansion:

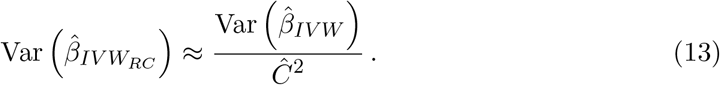

We note that this re-calibration is conceptually similar to MRlap when using nonoverlapping samples, although different approaches are used to estimate the calibration parameter, as discussed in Supplementary C.. While MRlap relies on strong assumptions about the underlying genetic architecture to estimate this parameter, regression calibration only uses data from the replication sample.

#### 2.2.1 Robust regression calibration

The IVW estimator using regression calibration relies on exactly the same assumptions as the standard IVW estimator and would only provide consistent estimates if the InSIDE assumption holds. In presence of correlated pleiotropy, such as in the model presented in (4) and (5), the IVW estimate using regression calibration 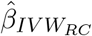 would be biased. In order to extend regression calibration to settings where the InSIDE assumption doesn’t hold, we could instead invoke the ‘majority’ rule on the set of pleiotropic effects *α*_*j*_s. That is, we assume that a majority of instruments have a zero pleiotropic effect. Assuming that genetic variants are ordered according to their ratio estimates, the simple median-based estimate [25] would be:

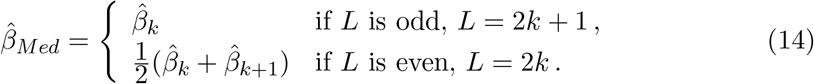

Using the same logic, it is possible to obtain a *Ĉ*_*M ed*_ corresponding to the median ratio 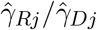 and to re-calibrate the causal effect as:

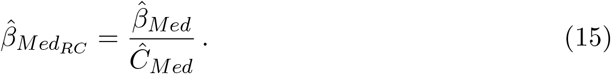

In fact, the regression calibration framework can be used to re-calibrate *any* pleiotropy robust two-sample MR estimator. For the purposes of this paper, we propose to use it with:

- The IVW estimator (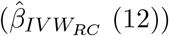) which relies on balanced pleiotropy;
- The simple median-based estimator (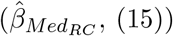), and the weighted median-based estimator 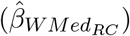 which rely on variations of the ‘majority’ rule;
- The simple mode-based estimator 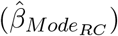 and the weighted mode-based estimator 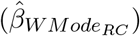 that rely on the zero modal pleiotropy assumption, or ‘plurality’ rule [6]

We use the standard implementation of all these estimators from the `twoSampleMR` R-Package [22] to obtain the causal effect estimates and the calibration parameters. All regression calibration estimators are implemented in the `RegressionCalibration` R-package to facilitate their use.

### 2.3 Simulation studies

We simulate SNP-exposure and SNP-outcome associations for *M* (30, 000) independent genetic variants, using data generating models (1) and (2). In all simulations, the causal effect is set to *β* = 0.2. We assume a spike-and-slab distribution for the SNP effect sizes, with a proportion *π*_*x*_ (0.1) of the genetic variants having a direct effect on the exposure, explaining a proportion 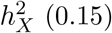 of the phenotypic variance, corresponding to the heritability of the exposure. Similarly, we assume that a proportion *π*_*y*_ of the genetic variants has a direct effect on the outcome, with a heritability of 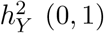. This corresponds to a scenario of balanced pleiotropy. In most simulations, the sample sizes for discovery, replication and outcome samples are respectively *N*_*D*_ = 90, 000, *N*_*R*_ = 50, 000, and *N*_*Y*_ = 50, 000, but we also simulate data using different ratios for discovery/replication (*N*_*D*_*/N*_*R*_), different total exposure sample size *N*_*D*_ + *N*_*R*_, and outcome sample size (*N*_*Y*_).

In addition to the balanced pleiotropy scenario, we simulate data in presence of a genetic confounder that induces correlated pleiotropy: Specifically, we explored the effect of weak (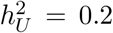, *π*_*U*_ = 0.01, *q*_*X*_ = 0.4, *q*_*Y*_ = 0.3) and strong (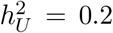, *π*_*U*_ = 0.0005, *q*_*X*_ = 0.4, *q*_*Y*_ = 0.3) correlation.

For each simulation scenario, we simulate 1,000 sets of SNP-exposure and SNP-outcome associations, and instruments for analyses are selected for different thresholds (*T*). In general, the genome-wide significance threshold (*T* ≈ 5.45) is used, either in the meta-analysis of discovery and replication results or in the discovery results (depending on the approach used) to select instruments to estimate the causal effect, but other thresholds were also used, to vary the strength of the instruments. For the balanced pleiotropy scenario, we use the following estimation strategies for the causal effect: the naive IVW; IVW using a 3-sample design; MR-RAPS using a 3-sample design; MR-RAPS using ZP; MR-RAPS using UMVCUE; and IVW using regression calibration approach. In addition, for the weak and strong correlated pleiotropy scenarios, we use the (simple and weighted) median- and mode-based implementations of regression calibration. For each scenario, we report the mean causal effect estimate, its standard deviation, as well as the bias and the root mean square error (RMSE).

### 2.4 Applied examples

To further compare the different approaches on real data, we adopt the strategy of Zhao et al. [10] of performing a ‘same-trait’ analyses to estimate the causal effect of body mass index (BMI) on itself, where we helpfully know the effect equals 1. We also assess the causal effect of BMI on systolic blood pressure (SBP). We use UK Biobank (UKB) [19] to create three independent samples of equal sample size (*N* = 126, 510), using only unrelated individuals of British ancestry (identified using genomic principal components). The first two samples are used for the exposure (BMI) and the third sample is used for the outcome (BMI and SBP). Phenotypic data is normalised (inverse-normal quantile transformed) and adjusted for the following covariates: sex, age, age^2^, and the first 40 principal components. GWAS is performed for each sample using BGENIE [26], using HapMap SNPs HapMap3 genetic variants [27] (*M* ≈ 1, 150, 000), keeping only SNPs with info quality *>* 0.9 and minor allele frequency *>* 0.01.

We repeat this process to generate GWAS summary statistics 100 times. For each repetition, causal effect estimates are estimated using all the approaches described above. Independent instruments for different p-value thresholds are selected from these summary statistics using the ‘clump_data()’ (default parameters, ‘clump_kb’ = 10,000, ‘clump_r2’ = 0.001,) from the ‘twoSampleMR’ R-package [22]. Note that in some cases, we also use random p-values (instead of observed ones) to select instruments (referred to as random LD-clumping) to avoid using the most strongly associated SNP in each region.

## 3 Results

### 3.1 Simulation studies

#### 3.1.1 Balanced pleiotropy

Table 1 and Figure 3 show results for the causal effect estimates using the battery of methods described. The naive IVW estimator is the most precise because it uses both discovery and replication data to identify instruments meeting the significance threshold. For this reason, a larger set of instruments (159 instead of 56 on average) are utilised and the naive IVW estimator has a lower RMSE than most estimators, but it is also the most biased. The 3-sample IVW approach strongly reduces the bias by removing Winner’s curse, but its estimate is consequently affected by weak-instruments bias. All other methods yield unbiased estimates by adjusting for weak instrument bias and Winner’s curse (either separately or jointly). MR-RAPS using ZP Winner’s curse correction is more precise than other unbiased approaches and yields the lowest RMSE, as it also uses combined data for the exposure.

**Table 1:** Mean, SD, bias, and RMSE of the different estimators in presence of balanced pleiotropy (1,000 simulations). Instruments are selected using a p-value threshold of 5 × 10^−8^ (T ≈ 5.45) and the true causal effect is 0.2. Using discovery data, an average of 57 instruments are used (F_D_ = 38.71, F_R_ = 15.44) whereas using combined data, an average of 159 instruments are used (F_C_ = 42.64).

| Estimator | Mean | SD | bias | RMSE |
| --- | --- | --- | --- | --- |
| Naive IVW | 0.175 | 0.0217 | - 0.0249 | 0.0330 |
| IVW, 3-sample design | 0.189 | 0.0347 | - 0.0107 | 0.0363 |
| MR-RAPS, 3-sample design | 0.202 | 0.0378 | 0.00164 | 0.0377 |
| MR-RAPS, ZP | 0.202 | 0.0268 | 0.00241 | 0.0269 |
| MR-RAPS, UMVCUE | 0.202 | 0.0370 | 0.00164 | 0.0371 |
| IVW, regression calibration | 0.203 | 0.0363 | 0.00280 | 0.0364 |

**Figure 3:**
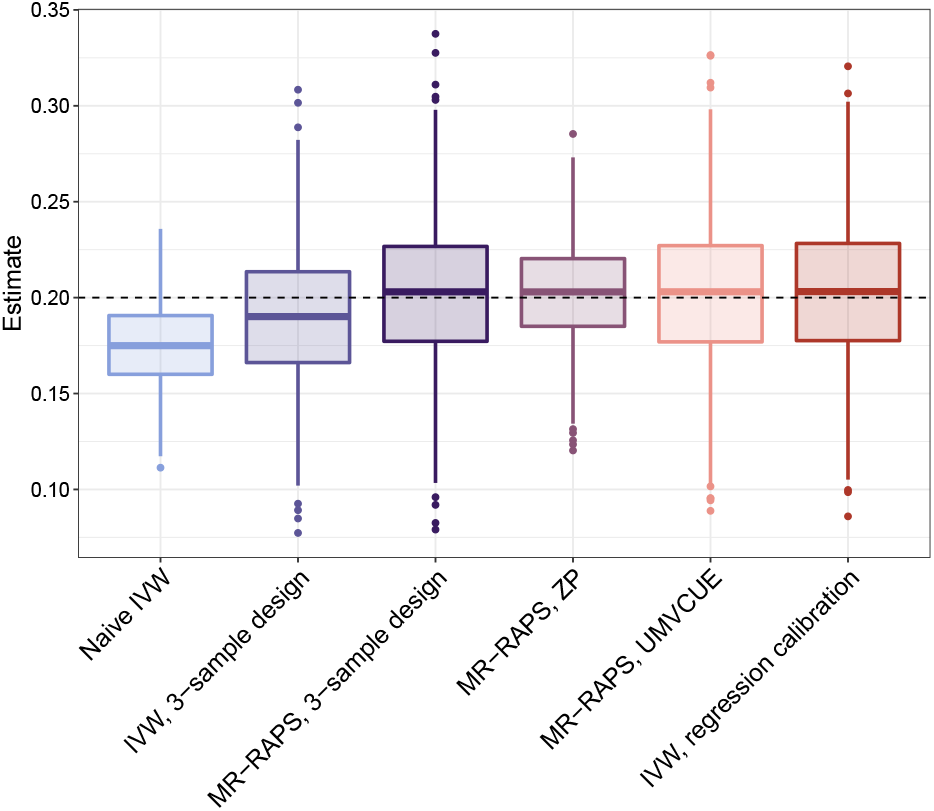
Distribution of estimates for naive IVW (light blue), IVW using a 3-sample design (dark blue), MR-RAPS using a 3-sample design (dark purple), MR-RAPS using ZP (light purple), MR-RAPS using UMVCUE (salmon pink) and IVW using the regression calibration approach (brick red) in presence of balanced pleiotropy (1,000 simulations). Instruments are selected using a p-value threshold of 5×10^−8^ (T ≈ 5.45) and the true causal effect (0.2) is indicated by the dashed line.

Varying the selection threshold does not affect the extent of the bias of the results for the 3-sample MR-RAPS approach, MR-RAPS using ZP, MR-RAPS using UMVCUE, and regression calibration (Figure 4 and Table S2). However, as expected, for methods that do not account for weak instruments (naive IVW and 3-sample IVW) the bias is stronger when less stringent selection thresholds are used. For naive IVW, when instruments are strong enough (*T >* 4.25, F-statistic *>* 18) the estimate reaches a plateau and the bias no longer appears to depend on the selection threshold. This is caused by two competing effects approximately cancelling out: increasing the threshold reduces weak-instruments bias, but it also increases Winner’s curse. Interestingly, when using less stringent thresholds, as the naive IVW estimator becomes more severely affected by weak instruments, robust approaches have lower RMSE despite their lower precision. For example, when *T* = 4.5, the RMSE for the naive IVW estimator is 0.0303, larger than the RMSE of any of the other estimators at this threshold.

**Figure 4:**
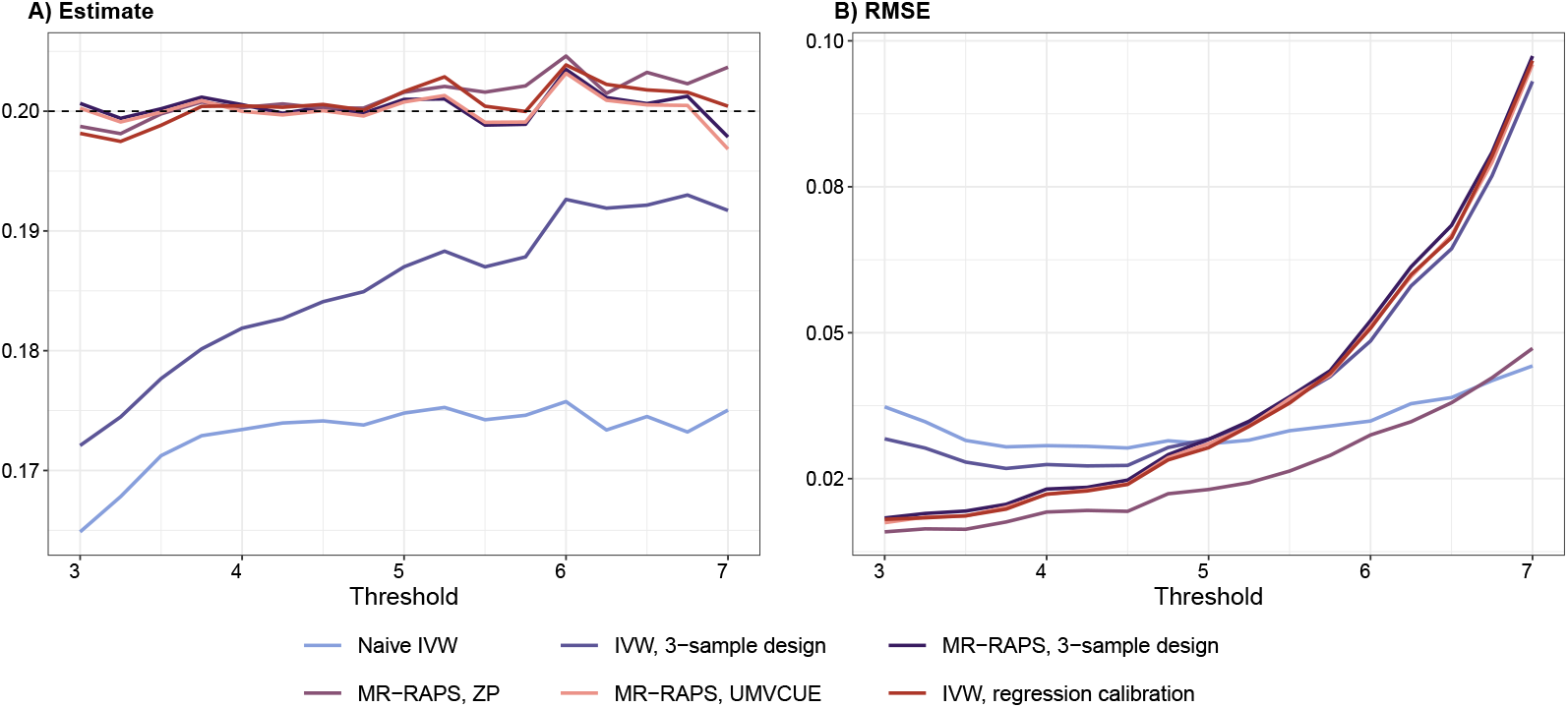
Effect of the selection threshold on naive IVW (light blue), IVW using a 3-sample design (dark blue), MR-RAPS using a 3-sample design (dark purple), MR-RAPS using ZP (light purple), MR-RAPS using UMVCUE (salmon pink) and IVW using the regression calibration approach (brick red) in presence of balanced pleiotropy (mean effect, panel A) and RMSE, panel B), across 1,000 simulations). Instruments are selected with different selection thresholds (T) and the true causal effect (0.2) is indicated by the dashed line in panel A).

#### 3.1.2 Effect of discovery-replication data splitting

To assess whether the relative size of the discovery and replication samples affects the performance of the different approaches, we fix the total exposure sample size (140,000) while varying the proportion that is used for discovery (from 35% up to 90%) (Figure 5 and Table S3). As expected, the naive IVW approach that relies on meta-analysed effects is not affected by varying the proportion since only the total exposure sample size matters. For all other methods, estimates are more precise when a larger proportion is used for discovery, as this corresponds to a larger set of instruments used. However, IVW using a 3-sample design is less biased when a smaller proportion of the sample is used for discovery, as this also reduces weak-instruments bias, and yields strongly biased estimates when the proportion gets higher (*>* 50%). In this case, the bias is even stronger than the naive IVW bias when the proportion used for discovery is larger than 70% and it seems preferable to use only about half of the sample for discovery, despite the lower precision. The 3-sample MR-RAPS, MR-RAPS using ZP, MR-RAPS using UMVCUE, and regression calibration are not affected by the proportion used and therefore for these methods, it is preferable to use a larger discovery sample, to increase precision. For example, when the proportion used for discovery is very large (≤ 85%), the naive IVW estimator and the IVW estimator using a 3-sample design have the largest RMSE, suggesting that the way of splitting the exposure data between discovery and replication can affect the bias-variance trade-off.

**Figure 5:**
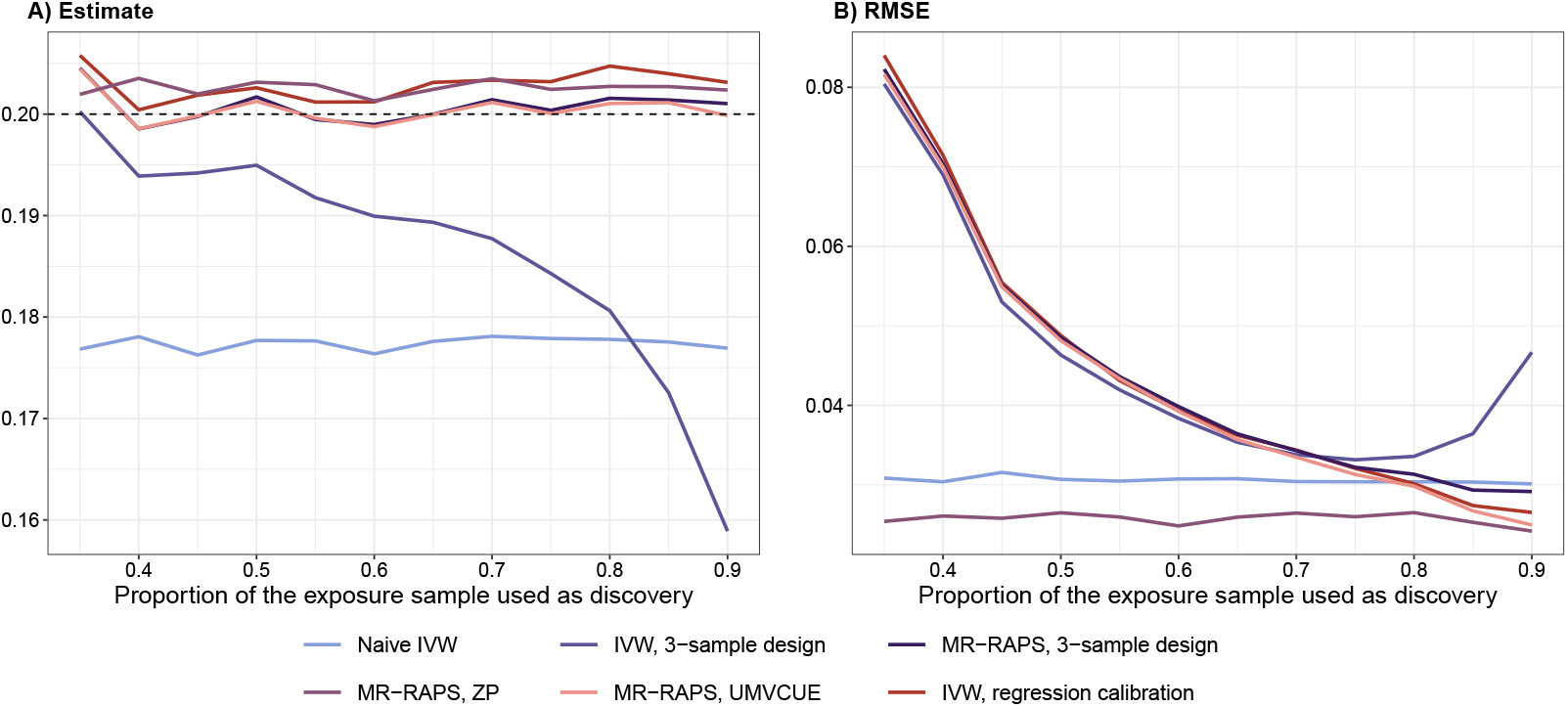
Effect of the proportion of the total exposure sample size used for discovery on naive IVW (light blue), IVW using a 3-sample design (dark blue), MR-RAPS using a 3-sample design (dark purple), MR-RAPS using ZP (light purple), MR-RAPS using UMVCUE (salmon pink) and IVW using the regression calibration approach (brick red) in presence of balanced pleiotropy (mean effect, panel A) and RMSE, panel B), across 1,000 simulations). Instruments are selected using a p-value threshold of 5 × 10^−8^ (T ≈ 5.45) and the true causal effect (0.2) is indicated by the dashed line in panel A).

We also performed simulations varying the total exposure sample size but keeping the proportion used for discovery and replication constant. Varying the total exposure sample size only affects estimates from methods that are susceptible to weak instruments (naive IVW and 3-sample IVW) and all other methods perform similarly for a wide range of sample sizes, with greater precision for larger sample sizes (Figure S1 and Table S4). For extremely large sample sizes (*N* ≤ 280, 000), the increase of precision from using the combined data is less important and the RMSE of the naïve IVW estimator is larger than the RMSE of the other estimators. When increasing the outcome sample size, approaches that correct for Winner’s curse and weak instruments have lower RMSE and a better bias-variance trade-off (Figure S2 and Table S5).

#### 3.1.3 Correlated Pleiotropy violating InSIDE

Finally, simulations are performed assuming correlated pleiotropy induced via two mechanisms: firstly, we model weak correlated pleiotropic effects across a large number of SNPs that influence a confounder (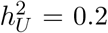, *π*_*U*_ = 0.01, *q*_*X*_ = 0.4, *q*_*Y*_ = 0.3) and secondly, we model strong pleiotropic effects induced by a smaller number of SNPs that influence the confounder (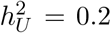, *π*_*U*_ = 0.0005, *q*_*X*_ = 0.4, *q*_*Y*_ = 0.3). Naively using SNPs with correlated pleiotropic effects as valid instruments in either an IVW or MR-RAPS analysis will generally lead to substantial bias. Indeed, if some SNPs are associated with the exposure *only* through the confounder, their ratio estimate would reflect the ratio of the effects of the confounder on both traits (e.g. *q*_*Y*_ */q*_*X*_ = 0.75) rather than the true underlying causal effect (*β* = 0.2). Moreover, the process of instrument selection can act to increase the extent of correlated pleiotropy depending on the threshold used, because it can enrich the included instrument set with strong and highly pleiotropic instruments. Such SNPs get a large weight in the analysis but also contain the most bias. It is in this setting where we envisage robust regression calibration based on median- or mode-based estimation being most useful because they are robust to correlated pleiotropy providing their respective majority or plurality assumptions hold. The results of weak and strong correlated pleiotropy scenarios are respectively shown in Table 2 and Figure 6).

**Table 2:**
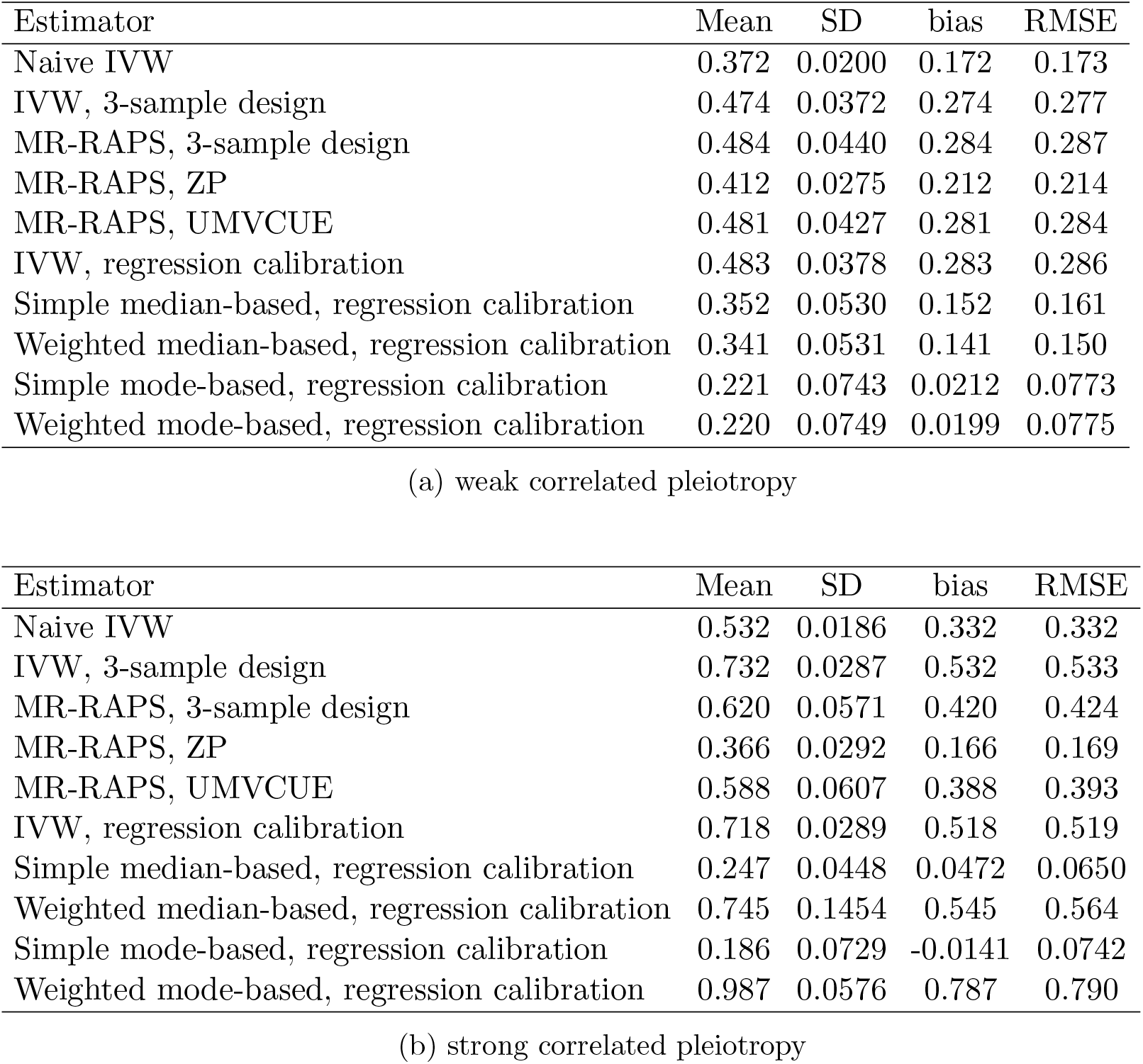
Mean, SD, bias, and RMSE of the different estimators in presence of correlated pleiotropy. Instruments were selected using a p-value threshold of 5 × 10^−8^ (T ≈ 5.45) and the true causal effect is 0.2. In presence of weak correlated pleiotropy (a), using discovery data, an average of 84 instruments are used (F_D_ = 41.92, F_R_ = 18.21, 32.7% of invalid instruments) whereas using combined data, an average of 206 instruments are used (F_C_ = 46.07, 23.6% of invalid instruments). In presence of strong correlated pleiotropy (b), using discovery data, an average of 66 instruments are used (F_D_ = 85.15, F_R_ = 42.05, 14.4% of invalid instruments) whereas using combined data, an average of 169 instruments are used (F_C_ = 71.65, 6.4% of invalid instruments).

**Figure 6:**
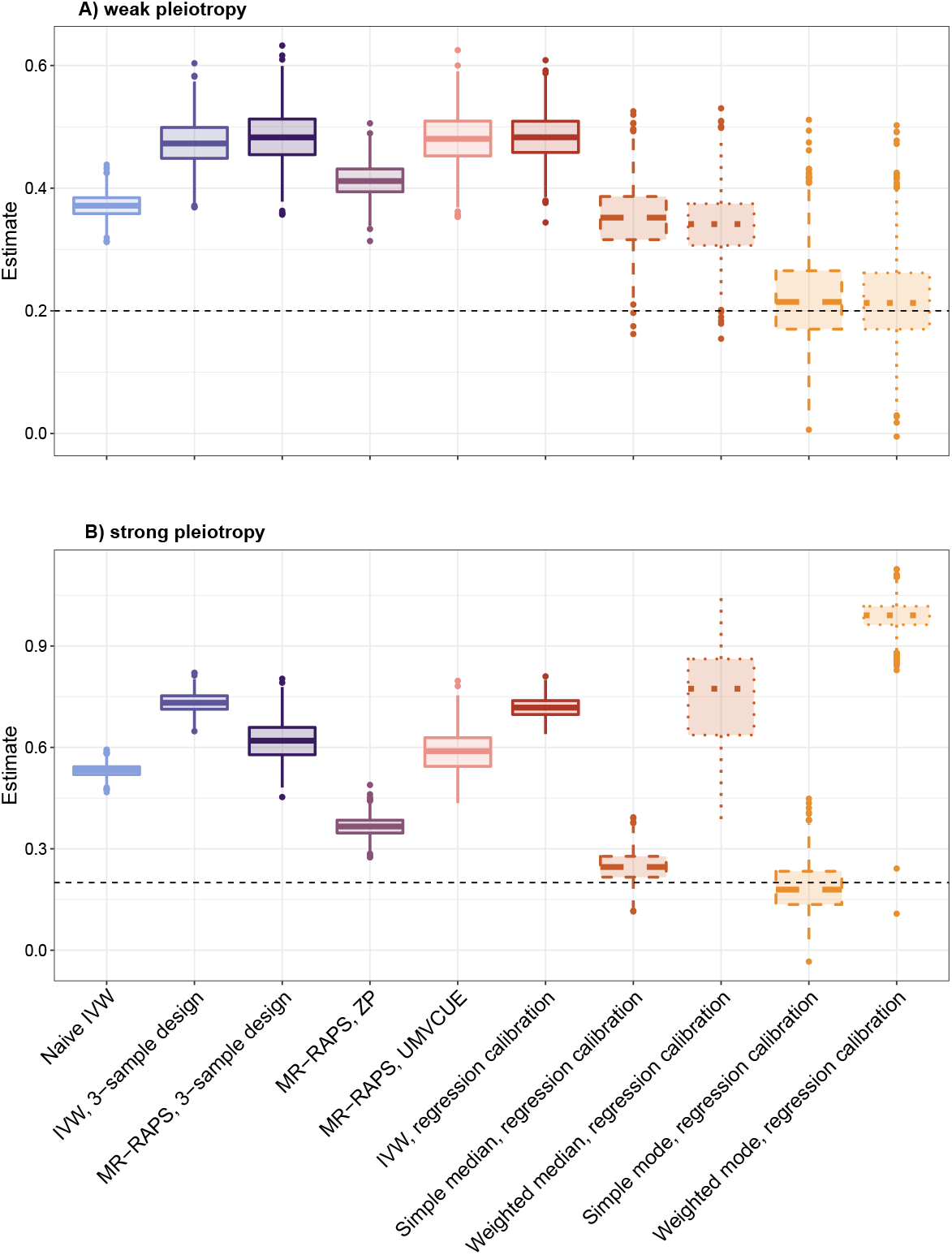
Distribution of estimates for naive IVW (light blue), IVW using a 3-sample design (dark blue), MR-RAPS using a 3-sample design (dark purple), MR-RAPS using ZP (light purple), MR-RAPS using UMVCUE (salmon pink), IVW using the regression calibration approach (brick red), simple median-based using the regression calibration approach (dark orange, dashed), weighted median-based using the regression calibration approach (dark orange, dotted), simple mode-based using the regression calibration approach (light orange, dashed), weighted mode-based using the regression calibration approach (light orange, dotted) in presence of weak (Panel A) and strong (Panel B) correlated pleiotropy (1,000 simulations). Instruments were selected using a p-value threshold of 5 × 10^−8^ (T ≈ 5.45) and the true causal effect (0.2) is indicated by the dashed line.

As expected, most methods (naive IVW, 3-sample IVW, 3-sample MR-RAPS, MR-RAPS ZP and MR-RAPS UMVCUE as well as regression calibration IVW) are strongly affected by the pleiotropic effects. Naive IVW appears to be less biased because of the different biases acting in opposite directions (bias towards the null from Winner’s curse and weak instruments bias upwards from correlated pleiotropy). Estimates from the median- and mode-based regression calibration approaches were the least precise, but also least biased when their assumptions were met and exhibited lower RMSE. However, this was highly dependent on the type of correlated pleiotropy simulated. In presence of weak but widespread correlated pleiotropy, the mode-based estimators perform better than the median-based estimators, and when only a small number of instruments with strong effects were responsible for the correlation, the ‘weighted’ approaches yielded highly biased results.

As expected, when the selection threshold used for instrument discovery increases, more instruments violating the independence assumption are included, leading to stronger bias (Figure S3, Tables S6 and S7). For instance, in the weak correlated pleiotropy scenario, when *T* = 5, it corresponds to 22% - 29% of invalid instruments (using combined and using discovery samples, respectively) whereas when *T* = 6, the proportion of invalid instruments goes up to 30% - 44%. The regression calibration approach using median-based estimators becomes more severely biased when *T >* 6.5, as the proportion of invalid instruments in the discovery samples goes over 50%. In the presence of strong correlated pleiotropy, the simple median-based and simple mode-based estimators lead to unbiased estimates for most selection thresholds and only exhibit bias when a larger proportion of instruments are invalid (when *T >* 7.5, more than 30% of instruments are invalid). Despite having a larger variance than other estimators, their RMSE was lower than the one of the naive IVW for all selection thresholds.

### 3.2 Applied examples

#### 3.2.1 Same trait analysis: Body Mass Index

To further investigate the potential of the different methods when applied to real data, we first focus on the causal effect of Body Mass Index (BMI) on itself. We estimate the causal effect of BMI on itself using various approaches and repeated the analysis in 100 random samples taken from UKB. We can see that naive IVW, while using more instruments than other methods (172 vs 55) is strongly biased towards the null 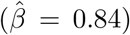 (Figure 7 A) and Table 3), mainly because of Winner’s curse since instruments are quite strong (F-statistic = 55.17). Its estimate is incompatible with a causal effect of 1 (95% confidence interval: 0.810-0.869). All other methods seem to be able to recover the true expected causal effect of 1, with the exception of MR-RAPS using ZP. This differs from the simulation results, for which this approach yielded unbiased estimates. We hypothesise that this could be caused by a failure to adequately model the LD structure in the data, which would mean that the estimated bias correction made for Winner’s curse for each SNP ranking top in a specific gene region is not correct. We explain this further in Supplementary D.. Indeed, when applying the MR-RAPS using ZP approach with a random SNP in each region, the mean absolute bias was lower (0.059 vs 0.029) but this comes at the cost of a lower precision (Figure S4 and Table S8), and to a lesser extent, MR-RAPS using UMVCUE is also affected by this issue.

**Figure 7:**
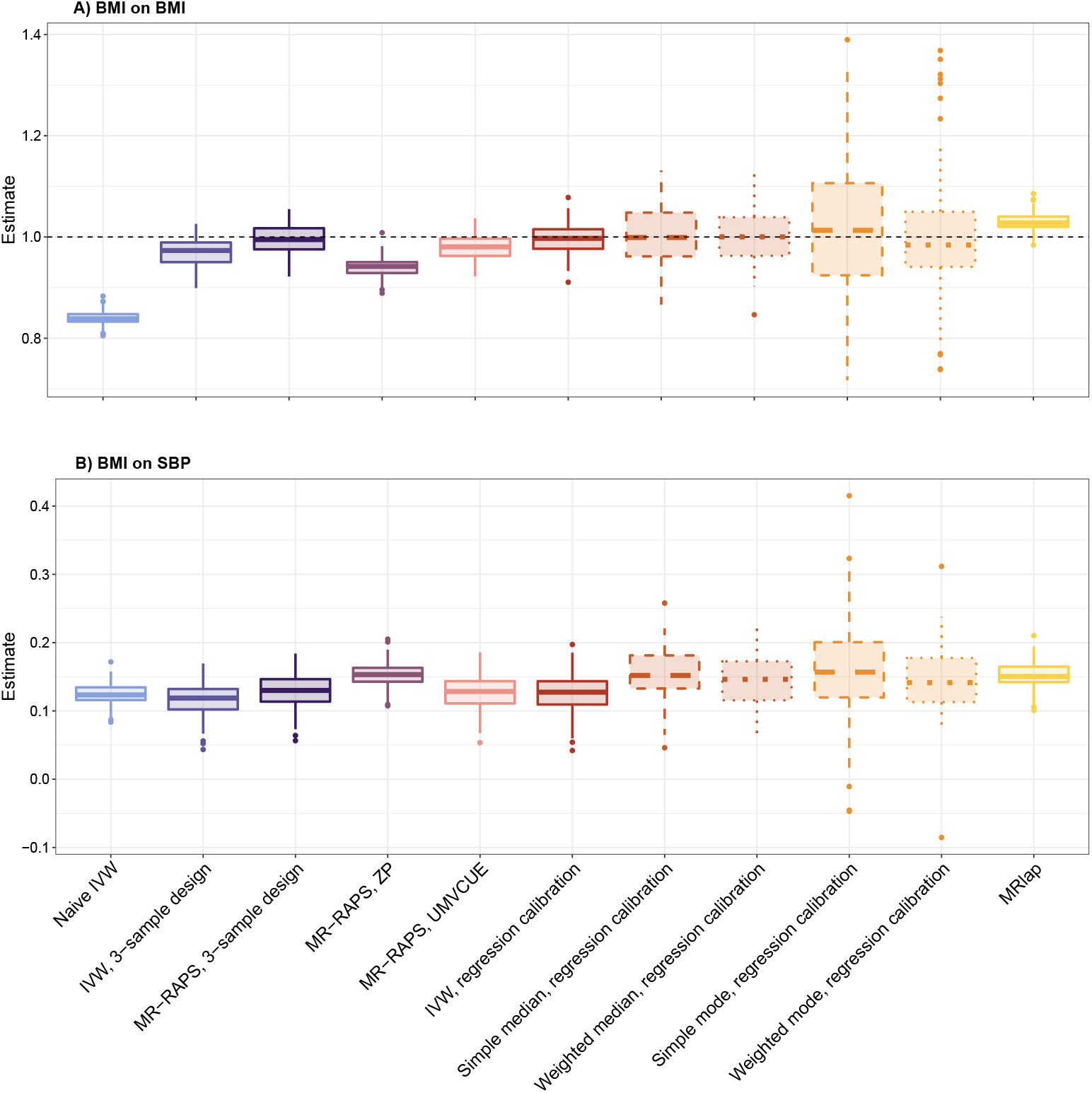
Distribution of causal effect estimates of BMI on BMI, Panel A) (expected to be 1, dashed line) and of BMI on SBP, Panel B), using naive IVW (light blue), IVW using a 3-sample design (dark blue), MR-RAPS using a 3-sample design (dark purple), MR-RAPS using ZP (light purple), MR-RAPS using UMVCUE (salmon pink), IVW using the regression calibration approach (brick red), simple median-based using the regression calibration approach (dark orange, dashed), weighted median-based using the regression calibration approach (dark orange, dotted), simple mode-based using the regression calibration approach (light orange, dashed), weighted mode-based using the regression calibration approach (light orange, dotted) and MRlap (yellow) (from 100 random samplings). Instruments were selected using a p-value threshold of 5 × 10^−8^ (T ≈ 5.45).

**Table 3:**
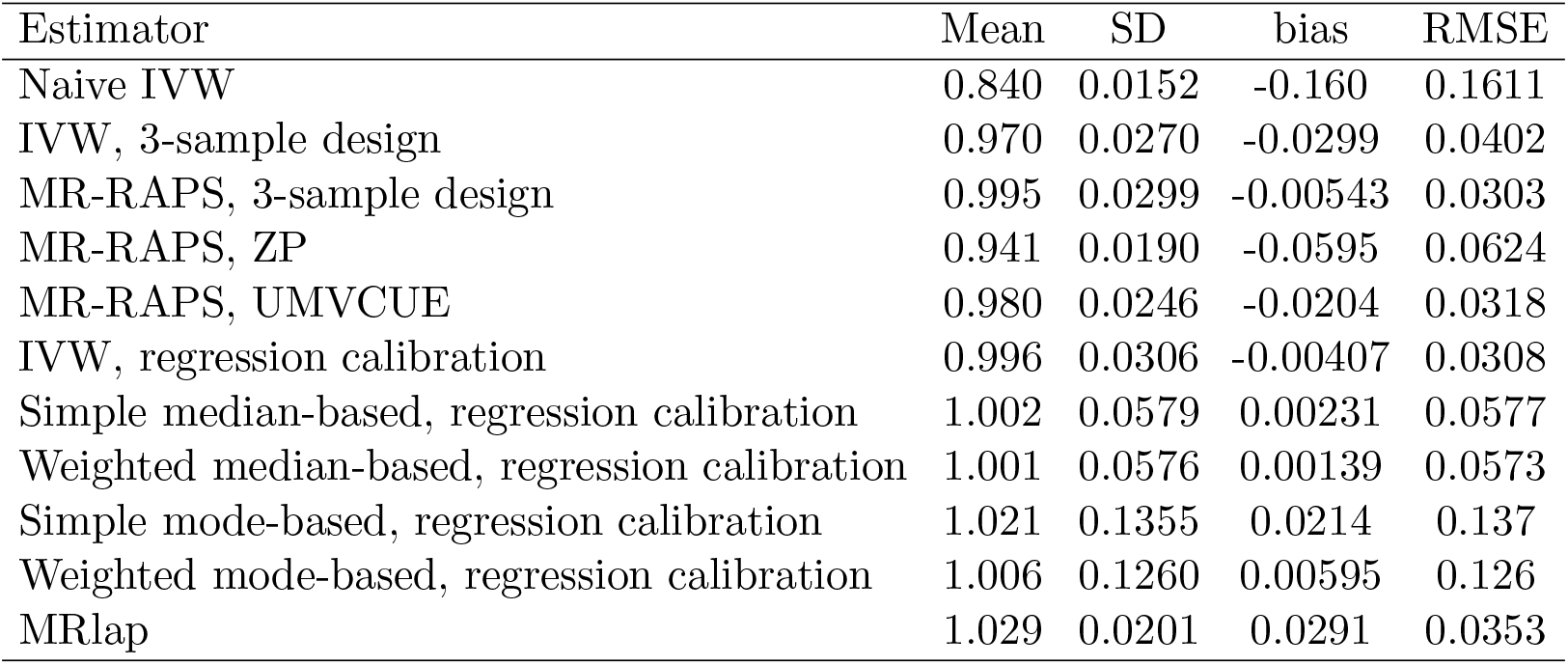
Mean, SD, bias, and RMSE of the different estimators when estimating the causal effect of BMI on itself. Instruments were selected using a p-value threshold of 5 × 10^−8^ (T ≈ 5.45) and the true causal effect is expected to be 1. Using discovery data, an average of 55 instruments are used (F_D_ = 51.26, F_R_ = 37.64) whereas using combined data, an average of 172 instruments are used (F_C_ = 55.17).

| Estimator | Mean | SD | bias | RMSE |
| --- | --- | --- | --- | --- |
| Naive IVW | 0.840 | 0.0152 | -0.160 | 0.1611 |
| IVW, 3-sample design | 0.970 | 0.0270 | -0.0299 | 0.0402 |
| MR-RAPS, 3-sample design | 0.995 | 0.0299 | -0.00543 | 0.0303 |
| MR-RAPS, ZP | 0.941 | 0.0190 | -0.0595 | 0.0624 |
| MR-RAPS, UMVCUE | 0.980 | 0.0246 | -0.0204 | 0.0318 |
| IVW, regression calibration | 0.996 | 0.0306 | -0.00407 | 0.0308 |
| Simple median-based, regression calibration | 1.002 | 0.0579 | 0.00231 | 0.0577 |
| Weighted median-based, regression calibration | 1.001 | 0.0576 | 0.00139 | 0.0573 |
| Simple mode-based, regression calibration | 1.021 | 0.1355 | 0.0214 | 0.137 |
| Weighted mode-based, regression calibration | 1.006 | 0.1260 | 0.00595 | 0.126 |
| MRlap | 1.029 | 0.0201 | 0.0291 | 0.0353 |

Mostly, approaches differ regarding their precision: the mode-based estimators are the least precise whereas MR-RAPS using ZP and MRlap are most precise after naive IVW (as they also use combined data for the exposure), followed by MR-RAPS using UMVCUE, as expected since the UMVCUE estimator provides the most efficient way of combining discovery and replication data. MRlap yields a slight upward bias (possibly due to a violation of the spike-and-slab genetic architecture assumption needed by the approach), with the estimates still being compatible with a causal effect of 1 and an absolute bias much lower than the one from naive IVW. The naive IVW estimator has the worst RMSE, which is 5 times larger than the RMSE of the approaches that account for Winner’s curse and weak instrument bias.

#### 3.2.2 Effect of Body Mass Index on Systolic Blood Pressure

We use the same sampling approach (100 times 3 non-overlapping samples from UKB) to estimate the causal effect of BMI on systolic blood pressure (SBP) using various methods. When selecting instruments using a p-value threshold of 5 *×* 10^−8^, surprisingly, the naive IVW estimate is not the weakest one, as it should be because of weak instruments and Winner’s curse biasing the estimate towards the null (Figure 7 B) and Table 4). This could be due to heterogeneity in the causal effect: using the naive IVW approach and combining discovery and replication data permits the use of a much larger number of instruments (172 vs 55) than most other approaches (MR-RAPS using ZP and MRlap being the only other ones using the combined data to select instruments). If there exist multiple underlying causal mechanisms, it is possible some of the instruments would act through different mechanisms, and depending on the set of instruments used, the causal effect estimate might vary accordingly.

**Table 4:** Mean and SD of the different estimators when estimating the causal effect of BMI on SBP. Instruments were selected using a p-value threshold of 5 × 10^−8^ (T ≈ 5.45). Using discovery data, an average of 55 instruments are used (F_D_ = 51.26, F_R_ = 37.64) whereas using combined data, an average of 172 instruments are used (F_C_ = 55.17).

| Estimator | Mean | SD |
| --- | --- | --- |
| Naive IVW | 0.125 | 0.0163 |
| IVW, 3-sample design | 0.116 | 0.0249 |
| MR-RAPS, 3-sample design | 0.129 | 0.0260 |
| MR-RAPS, ZP | 0.153 | 0.0179 |
| MR-RAPS, UMVCUE | 0.127 | 0.0262 |
| IVW, regression calibration | 0.126 | 0.0280 |
| Simple median-based, regression calibration | 0.154 | 0.0376 |
| Weighted median-based, regression calibration | 0.144 | 0.0359 |
| Simple mode-based, regression calibration | 0.161 | 0.0755 |
| Weighted mode-based, regression calibration | 0.146 | 0.0504 |
| MRlap | 0.153 | 0.0203 |

To further explore this hypothesis, we focus only on a subset of methods that used the same set of instruments (IVW using a 3-sample design, IVW using the regression calibration approach, and median-based and mode-based approaches) and compare causal effect estimates for different selection thresholds, i.e. different subsets of instruments (Figure 8 and Table S9). If the homogeneity of the causal effect assumption was holding, we would expect the estimate from IVW using a 3-sample design to be more strongly biased towards the null when using weaker instruments, and the estimates from the other approaches should not vary depending on the threshold. However, the causal effect estimates decrease when the threshold used to select instruments is more stringent. The ‘weighted’ (median-based and mode-based) approaches however seem to be able to provide consistent causal effect estimates, suggesting the fact that the majority of the instruments, in terms of weights, correspond to a single causal mechanism independently of the threshold used for selection.

**Figure 8:**
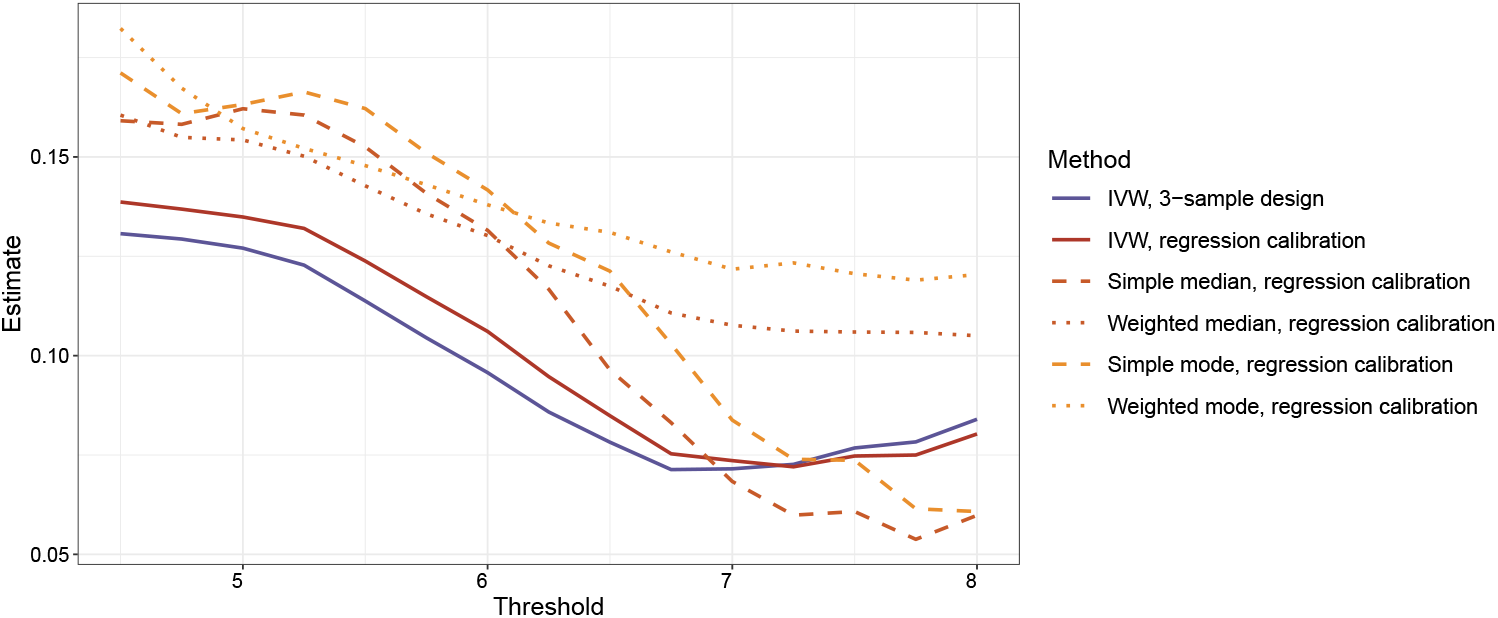
Effect of the selection threshold when estimating the causal effect of BMI on SBP on IVW using a 3-sample design (dark blue), IVW using the regression calibration approach (brick red), simple median-based using the regression calibration approach (dark orange, dashed), weighted median-based using the regression calibration approach (dark orange, dotted), simple mode-based using the regression calibration approach (light orange, dashed), weighted mode-based using the regression calibration approach (light orange, dotted). Instruments were selected with different selection thresholds (T).

## 4 Discussion

In this paper, we present regression calibration as a novel approach for summary data MR when discovery and replication samples are available for the exposure. We compare its performance to those of different approaches that can account for Winner’s curse and weak instruments bias. We first conduct simulation studies and observe that both Winner’s curse and weak instruments contribute to bias in two-sample MR causal effect estimates. However, when selecting IVs using genome-wide significance threshold (5*×*10^−8^), the extent of weak instrument bias is relatively small, and correcting for Winner’s curse using a 3-sample design can already strongly attenuate the bias.

When the InSIDE assumption holds, various approaches are able to accurately recover the true causal effect estimates. Amongst these approaches robust to both Winner’s curse and weak instrument bias, some do not explicitly rely on a discovery sample. This is the case for MR-RAPS using ZP, which can use combined data (discovery+replication) for the exposure and therefore exhibit increased precision due to the use of a larger sample size for instrument selection. Other 3-sample approaches also yield unbiased estimates, with the UMVCUE approach being slightly more efficient. The bias-variance trade-off of all the approaches strongly depends on the discovery, replication, and outcome sample sizes. For larger - but not unrealistic nowadays - sample sizes, the trade-off is clearly in favour of using a 3-sample design to account for Winner’s curse and weak instrument bias, especially if this is possible to use a large discovery sample and a smaller replication sample. In this case, the gain in precision obtained by combining the discovery and the replication samples is not important enough to compensate for the bias induced by Winner’s curse and weak instruments using the naive IVW approach.

The regression calibration approach yields unbiased estimates in balanced pleiotropy settings as well as more complex (and arguably more realistic) correlated pleiotropy settings via median and mode-based implementations such. However, they lose precision compared to other approaches.

We also perform a compelling same-trait analysis, estimating the causal effect of BMI on itself. Since in this case, we know the true causal effect is one, it allows us to evaluate the strength of the bias for real data, where estimator performance was degraded to varying degrees by mis-specification of the genetic architecture, selection mechanism, and pleiotropy distribution. For example, MR-RAPS using ZP and to a lesser extent MR-RAPS using UMVCUE yields downward biased causal effect estimates. This is likely because of the LD-clumping step that is needed with real data to obtain independent instruments. Unless the top SNP in a specific region is independent of all other signals, the selection bias (and hence selection threshold) assumed by the ZP and UMVCUE corrections will be incorrect (see Figure S5). In the case of the UMVCUE, more sophisticated approaches do exist to account for LD structure [16] but are also prone to mis-specification. MRlap is also slightly biased, and this is probably because it makes the strongest parametric assumptions about the genetic architecture of the exposure. Violations of these assumptions could result into sub-optimal correction. Regression calibration avoids explicit modelling of the selection mechanism or genetic architecture and so is not affected by either issue, which is a real strength of the approach. Combined with its ability to deal with correlated pleiotropy, this robustness sets them apart from all other approaches. As future work we will look to extend the technique of regression calibration to more complex multi-parameter MR modelling frameworks, for example, cluster-analysis-based methods [20, 21] and non-linear MR [28] as well as to settings where sample overlap is allowed.

When assessing the causal effect of BMI on SBP, we observe evidence of heterogeneity in the causal effect. Indeed, even the causal effect estimates from approaches robust to Winner’s curse and weak instrument bias vary with the threshold, yielding weaker estimates for more stringent thresholds. This is not necessarily indicating the presence of correlated pleiotropy, and could also be due to distinct causal mechanisms underlying different subsets of genetic variants. Using regression calibration with weighted mode-based and median-based, results are more consistent and less strongly affected by the selection threshold, suggesting that the majority of the variants (in terms of weight) is capturing a stronger causal effect.

As illustrated in Sadreev et al [29], Winner’s curse leads to a substantial overestimation in SNP-trait associations and this has direct implications for MR studies. Further descriptive work focusing on a single relationship, BMI on coronary artery disease, suggests that Winner’s curse does not materially affect the highly statistically significant causal estimate [30]. This is not surprising as the deleterious effect of BMI on coronary artery disease is well established, but may not be the case for other relationships. Our results suggest that Winner’s curse can lead to strong bias, important under-coverage and could hinder the identification of modest causal effects. Hence, it should not be ignored and there exist various approaches to deal with it. Regression calibration has been implemented for several estimators in the `RegressionCalibration` R-package that works seamlessly with `TwoSampleMR`.

## Acknowledgements

We thank Lausanne University Hospital for use of their HPC cluster to perform the simulation study in this paper.

## Data availability

The genotype and phenotype data underlying the results presented in the Applied examples section are third-party data available via application to the UK Biobank. Access to data was obtained under application number 16389. The R-package providing the regression calibration estimators is available on GitHub (https://github.com/n-mounier/RegressionCalibration). Datasets for simulation studies can be generated through the code ‘`launch_simulations.R`’, and results can be reproduced using the code ‘`make_figures_simulations.R`’ from the GitHub repository.

## Author Contributions

NM: *Conceptualization, Data Curation, Formal Analysis, Methodology, Software, Writing – Original Draft Preparation, Writing – Review & Editing*

DSR: *Conceptualization, Data Curation, Methodology, Software, Writing – Original Draft Preparation, Writing – Review & Editing*

ZK: *Conceptualization, Methodology, Writing – Review & Editing*

FD: *Conceptualization, Methodology, Writing – Review & Editing*

JB: *Conceptualization, Methodology, Supervision, Writing – Original Draft Preparation, Writing – Review & Editing*

## References

[1] Smith GD, Ebrahim S. ‘Mendelian randomization’: can genetic epidemiology contribute to understanding environmental determinants of disease? International Journal of Epidemiology. 2003;32(1):1–22.

[2] Burgess S, Butterworth A, Thompson SG. Mendelian Randomization Analysis With Multiple Genetic Variants Using Summarized Data. Genet Epidemiol. 2013;37:658–65.

[3] Bowden J, Holmes MV. Meta-analysis and Mendelian randomization: A review. Research Synthesis Methods. 2019;10(4):486–96.

[4] Hemani G, Zheng J, Elsworth B, Wade KH, Haberland V, Baird D, et al. The MR-Base platform supports systematic causal inference across the human phenome. eLife. 2018;7:e34408.

[5] Bowden J, Del Greco M F, Minelli C, Davey Smith G, Sheehan N, Thompson J. A framework for the investigation of pleiotropy in two-sample summary data Mendelian randomization. Statistics in medicine. 2017;36(11):1783–802.

[6] Hartwig FP, Smith GD, Bowden J. Robust inference in summary data Mendelian randomization via the zero modal pleiotropy assumption. International Journal of Epidemiology. 2017;46(6):1985–98.

[7] Verbanck M, Chen CY, Neale B, Do R. Detection of widespread horizontal pleiotropy in causal relationships inferred from Mendelian randomization between complex traits and diseases. Nature Genetics. 2018;50(5):693–8.

[8] Bowden J, Del Greco M F, Minelli C, Davey Smith G, Sheehan NA, Thompson JR. Assessing the suitability of summary data for two-sample Mendelian randomization analyses using MR-Egger regression: the role of the I2 statistic. International journal of epidemiology. 2016;45(6):1961–74.

[9] Bowden J, Davey Smith G, Burgess S. Mendelian randomization with invalid instruments: effect estimation and bias detection through Egger regression. International journal of epidemiology. 2015;44(2):512–25.

[10] Zhao Q, Wang J, Hemani G, Bowden J, Small DS. Statistical inference in two-sample summary-data Mendelian randomization using robust adjusted profile score. The Annals of Statistics. 2020;48(3):1742–69.

[11] Bowden J, Spiller W, Del Greco M F, Sheehan N, Thompson J, Minelli C, et al. Improving the visualization, interpretation and analysis of two-sample summary data Mendelian randomization via the Radial plot and Radial regression. International Journal of Epidemiology. 2018;47(4):1264–78.

[12] Zhong H, Prentice RL. Bias-reduced estimators and confidence intervals for odds ratios in genome-wide association studies. Biostatistics. 2008;9(4):621–34.

[13] Xiao R, Boehnke M. Quantifying and correcting for the winner’s curse in genetic association studies. Genetic Epidemiology: The Official Publication of the International Genetic Epidemiology Society. 2009;33(5):453–62.

[14] Palmer C, Pe’er I. Statistical correction of the Winner’s Curse explains replication variability in quantitative trait genome-wide association studies. PLOS Genetics. 2017;13(7):e1006916.

[15] Bowden J, Dudbridge F. Unbiased estimation of odds ratios: combining genomewide association scans with replication studies. Genetic Epidemiology: The Official Publication of the International Genetic Epidemiology Society. 2009;33(5):406–18.

[16] Robertson DS, Prevost AT, Bowden J. Accounting for selection and correlation in the analysis of two-stage genome-wide association studies. Biostatistics. 2016;17(4):634–49.

[17] Carroll RJ, Stefanski LA. Approximate Quasi-likelihood Estimation in Models with Surrogate Predictors. Journal of the American Statistical Association. 1990;85(411):652–63.

[18] Spiegelman D, McDermott A, Rosner B. Regression calibration method for correcting measurement-error bias in nutritional epidemiology. The American Journal of Clinical Nutrition. 1997;65(4):1179S–1186S.

[19] Sudlow C, Gallacher J, Allen N, Beral V, Burton P, Danesh J, et al. UK Biobank: An Open Access Resource for Identifying the Causes of a Wide Range of Complex Diseases of Middle and Old Age. PLoS Med. 2015;12:1–10.

[20] Foley CN, Mason AM, Kirk PDW, Burgess S. MR-Clust: clustering of genetic variants in Mendelian randomization with similar causal estimates. Bioinformatics. 2020;37(4):531–41.

[21] Shapland CY, Zhao Q, Bowden J. Profile-likelihood Bayesian model averaging for two-sample summary data Mendelian randomization in the presence of horizontal pleiotropy. Statistics in Medicine. 2022;41(6):1100–19.

[22] Hemani G, Zheng J, Elsworth B, Wade K, Baird D, Haberland V, et al. The MR-Base platform supports systematic causal inference across the human phenome. eLife. 2018;7:e34408.

[23] Bowden J, Glimm E. Unbiased estimation of selected treatment means in two-stage trials. Biometrical Journal: Journal of Mathematical Methods in Biosciences. 2008;50(4):515–27.

[24] Mounier N, Kutalik Z. Bias correction for inverse variance weighting Mendelian randomization. bioRxiv [Preprint]. 2021. bioRxiv 2021.03.26.437168 [posted 2021 March 28; revised 2022 Sep 20; cited 2022 Jan 10]. Available from: https://doi.org/10.1101/2021.03.26.437168.

[25] Bowden J, Davey Smith G, Haycock PC, Burgess S. Consistent estimation in Mendelian randomization with some invalid instruments using a weighted median estimator. Genetic epidemiology. 2016;40(4):304–14.

[26] Bycroft C, Freeman C, Petkova D, Band G, Elliott LT, Sharp K, et al. The UK Biobank resource with deep phenotyping and genomic data. Nature. 2018;562:203–9.

[27] International HapMap 3 Consortium. Integrating common and rare genetic variation in diverse human populations. Nature. 2010;467:52–8.

[28] Staley JR, Burgess S. Semiparametric methods for estimation of a nonlinear exposure-outcome relationship using instrumental variables with application to Mendelian randomization. Genetic Epidemiology. 2017;41(4):341–52.

[29] Sadreev II, Elsworth BL, Mitchell RE, Paternoster L, Sanderson E, Davies NM, et al. Navigating sample overlap, winner’s curse and weak instrument bias in Mendelian randomization studies using the UK Biobank. medRxiv [Preprint]. 2021. medRxiv 2021.06.28.21259622 [posted 2021 July 01; cited 2022 Jan 10]. Available from: https://doi.org/10.1101/2021.06.28.21259622.

[30] Jiang T, Gill D, Butterworth AS, Burgess S. An empirical investigation into the impact of winner’s curse on estimates from Mendelian randomization. International Journal of Epidemiology. 2022.

