## Supplementary Material for "Incorporating discovery and replication GWAS into summary data Mendelian randomization studies: A review of current methods and a simple, general and powerful alternative"

### Supplementary Materials

A. The ZP approach

B. The UMVCUE approachs

C. Equivalence between Regression Calibration and MRlap

D. Instrument selection in presence of LD

References

Supplementary Tables

Supplementary Figures

### A. The ZP approach

One obvious strategy for dealing with Winner's curse is to maximise the (log-)likelihood of the combined SNP-exposure association estimates $\hat{\gamma}_{Cj}$ conditional on being selected (at a given threshold $T$), with respect to the unknown parameter $\gamma_j$. Assuming a two-sided test and a normal unconditional distribution for the estimate, this yields:

$$\log f(\gamma_j|\hat{\gamma}_{Cj}, \sigma_{XCj}) - \log\left[\Phi\left(\frac{\gamma_j}{\sigma_{XCj}} - T\right) + \Phi\left(-\frac{\gamma_j}{\sigma_{XDj}} - T\right)\right], \tag{S1}$$

where $f(\cdot)$ represents a normal pdf. This procedure yields a bias-corrected version of $\hat{\gamma}_{Cj}$ which can then be used in the MR analysis. This general approach has been suggested multiple times in the literature [1, 2, 3]. We refer to this as the ZP approach throughout the paper.

### B. The UMVCUE approach

Unbiased estimate of the SNP-exposure associations can be obtained using the technique of Rao Blackwellisation [4, 5, 6]. This method calculates the expected value of the unbiased replication estimate $\hat{\gamma}_{Rj}$, conditional on a complete sufficient statistic for $\gamma_j$ and conditional on selection (for a selection threshold $T$). The resulting quantity is known as the Uniformly Minimum Variance Conditionally Unbiased Estimator (UMVCUE), denoted by $\tilde{\gamma}_j$. For two-sided testing, the UMVCUE is given by

$$\tilde{\gamma}_j = \hat{\gamma}_{MLE,j} - \frac{\sigma^2_{XRj}}{\sqrt{\sigma^2_{XRj} + \sigma^2_{XDj}}} \frac{\phi(S_{1j}) - \phi(S_{2j})}{\Phi(S_{1j}) + \Phi(-S_{2j})}, \tag{S2}$$

where

$$\hat{\gamma}_{MLE,j} = \frac{\sigma^2_{XRj}\hat{\gamma}_{Dj} + \hat{\sigma}^2_{Dj}\hat{\gamma}_{Rj}}{\sigma^2_{XRj} + \hat{\sigma}^2_{Dj}}, \tag{S3}$$

$$S_{1j} = \frac{\sqrt{\sigma^2_{XRj} + \sigma^2_{XDj}}}{\sigma^2_{XDj}} \left(\hat{\gamma}_{MLE,j} - T\sigma_{XDj}\right), \tag{S4}$$

$$S_{2j} = \frac{\sqrt{\sigma^2_{XRj} + \sigma^2_{XDj}}}{\sigma^2_{XDj}} \left(\hat{\gamma}_{MLE,j} + T\sigma_{XDj}\right). \tag{S5}$$

To use the UMVCUE in two-sample MR analyses, we substitute $\tilde{\gamma}_j$ for $\hat{\gamma}_{Dj}$ and the UMVCUE variance $\sigma^2_{\tilde{\gamma}j}$ for $\sigma^2_{Xj}$.

The UMVCUE perfectly removes Winner's curse bias in the SNP-exposure association estimate, and it is guaranteed to have a smaller variance than the replication estimate alone. However, the expression for its variance $\sigma^2_{\tilde{\gamma}j}$ does not have a convenient closed-form solution. Following previous recommendations, we propose to approximate using a parametric bootstrap algorithm to approximate $\sigma^2_{\tilde{\gamma}j}$, given the observed value of the UMVCUE $\tilde{\gamma}^*_j$.

For two-sided testing, the bootstrap procedure is as follows:

1. Define $\pi_{j+} = \Phi(-T + \tilde{\gamma}^*_j/\sigma_{XDj})$ and $\pi_{j-} = \Phi(-T - \tilde{\gamma}^*_j/\sigma_{XDj})$. With probability $\frac{\pi_{j+}}{\pi_{j+}+\pi_{j-}}$, generate bootstrap data $\hat{\gamma}^{(B)}_{Dj} \sim N(\tilde{\gamma}^*_j, \sigma^2_{XDj})$ conditional on $\hat{\gamma}^{(B)}_{Dj} \geq T\sigma_{XDj}$, and with probability $\frac{\pi_{j-}}{\pi_{j+}+\pi_{j-}}$, generate bootstrap data $\hat{\gamma}^{(B)}_{Dj} \sim N(\tilde{\gamma}^*_j, \sigma^2_{XDj})$ conditional on $\hat{\gamma}^{(B)}_{Dj} \leq -T\sigma_{XDj}$;

2. Generate bootstrap data $\hat{\gamma}^{(B)}_{Rj} \sim N(\tilde{\gamma}^*_j, \sigma^2_{XRj})$;

3. Calculate the UMVCUE $\tilde{\gamma}^{(B)}_j$ using the bootstrap data;

4. Repeat steps 1–3 to obtain a distribution of $\tilde{\gamma}^{(B)}_j$ values, and use their sample variance as an estimate for $\sigma^2_{\tilde{\gamma}j}$.

### C. Equivalence between Regression Calibration and MRlap

To account for Winner's curse, Regression Calibration and MRlap [7] both propose a direct correction of the causal effect estimate, rather than a correction of the SNP-exposure associations. In practice, it can be shown that Regression Calibration is a special case of MRlap, when samples are non-overlapping.

For simplicity we will use a notation similar to the one from (S4) in Mounier et al [7]:

$$\begin{aligned} \text{SNP-exposure assoc}^n \text{ in discovery sample: } \hat{\gamma}_{Dj} &= \gamma_j + \epsilon_{Dj}\,, & \text{(S6)} \\ \text{SNP-exposure assoc}^n \text{ in outcome sample: } \hat{\gamma}_{Yj} &= \gamma_j + \epsilon_{Yj}\,, & \text{(S7)} \\ \text{SNP-outcome assoc}^n \text{ in outcome sample: } \hat{\Gamma}_j &= \beta\gamma_{Yj} + \alpha_j + \varepsilon_{Yj}\,. & \text{(S8)} \end{aligned}$$

Instruments are selected based on their SNP-exposure association in the discovery sample, such as $|\hat{\gamma}_{Dj}| \cdot \sqrt{N_D} > T$. By denoting $S_j := \left\{\epsilon_{D_j} > \frac{T}{\sqrt{N_D}} - \gamma_j\right\} \cup \left\{\epsilon_{D_j} < -\frac{T}{\sqrt{N_D}} - \gamma_j\right\}$, the IVW causal effect estimate from (9) changes to:

$$\hat{\beta}_{IVW} \approx \frac{\sum_{j=1}^{M}(\hat{\Gamma}_j\,|S_j\,) \cdot (\hat{\gamma}_{Dj}\,|S_j\,) \cdot Pr(S_j)}{\sum_{j=1}^{M}(\hat{\gamma}_{Dj}\,|S_j\,)^2 \cdot Pr(S_j)}\,. \tag{S9}$$

Note that while $L$ denoted the number of instruments, $M$ represents the number of genome-wide markers from which instruments are selected. The expectation of the causal effect estimate can be written as:

$$
\begin{aligned}
E\left[\widehat{\alpha}_{IVW}\right] &\approx \frac{\sum_{j=1}^{M} E\left[\Big(\big(\beta \cdot \gamma_j + \alpha_j\big) + \big(\beta \cdot \epsilon_{Yj} + \varepsilon_j \,|S_j\big)\Big) \cdot \Big(\gamma_j + \big(\epsilon_{Dj} \,|S_j\big)\Big)\right] \cdot Pr(S_j)}{\sum_{j=1}^{M} E\left[\Big(\gamma_j + \big(\epsilon_{Dj} \,|S_j\big)\Big)^2\right] \cdot Pr(S_j)} \\
&= \frac{\sum_{j=1}^{M} E\left[\Big(\beta \cdot \big(\hat{\gamma}_{Yj} \,|S_j\big) + \big(\alpha_j + \varepsilon_{Yj} \,|S_j\big)\Big) \cdot \big(\hat{\gamma}_{Dj} \,|S_j\big)\right] \cdot Pr(S_j)}{\sum_{j=1}^{M} E\left[\big(\hat{\gamma}_{Dj} \,|S_j\big)^2\right] \cdot Pr(S_j)} . \qquad \text{(S10)}
\end{aligned}
$$

When the discovery and the outcome samples do not overlap $E\big[(\alpha_j + \varepsilon_{Yj}) \cdot \hat{\gamma}_{Dj}\big] = 0$, hence it simplifies to

$$
\begin{aligned}
E\left[\widehat{\alpha}_{IVW}\right] &\approx \frac{\sum_{j=1}^{M} E\left[\beta \cdot \big(\hat{\gamma}_{Yj} \,|S_j\big) \cdot \big(\hat{\gamma}_{Dj} \,|S_j\big)\right] \cdot Pr(S_j)}{\sum_{j=1}^{M} E\left[\big(\hat{\gamma}_{Dj} \,|S_j\big)^2\right] \cdot Pr(S_j)} \\
&= \beta \cdot \frac{\sum_{j=1}^{M} E\left[\big(\hat{\gamma}_{Yj} \,|S_j\big) \cdot \big(\hat{\gamma}_{Dj} \,|S_j\big)\right] \cdot Pr(S_j)}{\sum_{j=1}^{M} E\left[\big(\hat{\gamma}_{Dj} \,|S_j\big)^2\right] \cdot Pr(S_j)} . \qquad \text{(S11)}
\end{aligned}
$$

An obvious estimator for the ratio on the r.h.s. is

$$
\frac{\sum_{j=1}^{M} \big(\hat{\gamma}_{Yj} \,|S_j\big) \cdot \big(\hat{\gamma}_{Dj}\big)}{\sum_{j=1}^{M} \big(\hat{\gamma}_{Dj} \,|S_j\big)^2} . \qquad \text{(S12)}
$$

In the case of Regression Calibration, this is estimated using SNP-exposure associations from the replication sample rather than from the outcome sample, substituting $\hat{\gamma}_{Rj}$ for $\hat{\gamma}_{Yj}$ and using the regression coefficient estimate for

$$
\big(\hat{\gamma}_{Rj} \,|S\big) \;\sim\; \big(\hat{\gamma}_{Dj} \,|S\big) .
$$

Therefore, for non-overlapping samples, the IVW calibration parameter is equivalent to the MRlap one. However, very different approaches are used to estimate this calibration parameter. Regression Calibration only uses data from the replication sample while MRlap relies on a more complex modelling of the underlying genetic architecture to estimate this parameter without having to use additional data.

### D. Instrument selection in presence of LD

Most MR approaches, and in particular, the ones presented in this paper, require the use of independent instruments. Genetic variants located in the same genomic region are not independent, and linkage disequilibrium (LD) is a term used to describe the correlation structure. Because of LD, GWAS results only permit the identification of associated regions and not directly causal variants. Therefore, when selecting instruments for MR analyses, an additional step of LD-clumping is needed to keep only the most significant genetic association in each region in order to work with a set of independent instruments.

When correcting for Winner's curse, most approaches that correct SNP-exposure associations for Winner's curse or model the selection process are not taking this step into account, considering that a single threshold is used for all instruments. This is the case for MR-RAPS using ZP, MR-RAPS using UMVCUE or MRlap for example. However, the LD-clumping step may affect the threshold in each region. By keeping only the most strongly associated variant, it artificially makes the threshold more stringent, as the instrument does not only need to pass the threshold but also needs to be the most significant in the region (see LocusZoom plot [8], in Figure S5). As a consequence, most approaches will consider that Winner's curse bias for SNP-exposure associations is negligible when instruments that are strongly associated with the exposure ($\frac{\hat{\gamma}_i}{N} >> T$). This is not true, because in reality Winner's curse will also influence *which* SNP is selected as an instrument in each region and we would expect an overestimation of the SNP-exposure association even for instruments further away from the selection threshold.

To test how the LD-clumping step can affect the correction, we used random p-values (instead of observed ones) to perform random LD-clumping and avoid using the most strongly associated variant in each region (Figure S4). Estimates for the causal effect of BMI on itself are strongly biased towards the null when using the standard LD-clumping approach, suggesting that not taking into account this additional selection step reduces the correction performance. Selecting instruments using the random LD-clumping approach yields very different results: the causal effect seems to be slightly overestimated but is now compatible with a causal effect of 1. Note that while random LD-clumping seems to improve the correction, this comes at the cost of a lower precision because the set of instruments used is weaker.

### References


[1] Zhong H, Prentice RL. Bias-reduced estimators and confidence intervals for odds ratios in genome-wide association studies. Biostatistics. 2008;9(4):621-34.

[2] Xiao R, Boehnke M. Quantifying and correcting for the winner's curse in genetic association studies. Genetic Epidemiology: The Official Publication of the International Genetic Epidemiology Society. 2009;33(5):453-62.

[3] Palmer C, Pe'er I. Statistical correction of the Winner's Curse explains replication variability in quantitative trait genome-wide association studies. PLOS Genetics. 2017;13(7):e1006916.

[4] Bowden J, Glimm E. Unbiased estimation of selected treatment means in two-stage trials. Biometrical Journal: Journal of Mathematical Methods in Biosciences. 2008;50(4):515-27.

[5] Bowden J, Dudbridge F. Unbiased estimation of odds ratios: combining genomewide association scans with replication studies. Genetic Epidemiology: The Official Publication of the International Genetic Epidemiology Society. 2009;33(5):406-18.

[6] Robertson DS, Prevost AT, Bowden J. Accounting for selection and correlation in the analysis of two-stage genome-wide association studies. Biostatistics. 2016;17(4):634-49.

[7] Mounier N, Kutalik Z. Bias correction for inverse variance weighting Mendelian randomization. bioRxiv. 2021. Available from: `https://doi.org/10.1101/2021.03.26.437168`.

[8] Boughton AP, Welch RP, Flickinger M, VandeHaar P, Taliun D, Abecasis GR, et al. LocusZoom.js: interactive and embeddable visualization of genetic association study results. Bioinformatics. 2021;37(18):3017-8.

### Supplementary Tables

S1 *Summary of the data used by each approach for SNP selection and SNP-exposure association estimation.* . . . . . . . . . . . . . . . . . . . . 8

S8 *Mean absolute bias for naive IVW, MR-RAPS using ZP and MR-RAPS using UMVCUE when estimating the causal effect of BMI on itself, comparing two LD-clumping approaches. Instruments were selected using a p-value threshold of* $5 \times 10^{-8}$ *(*$T \approx 5.45$*). Using discovery data, an average of 55 instruments are used (*$F_D = 51.26$*,* $F_R = 37.64$*) using standard LD-clumping whereas an average of 55 instruments are used (*$F_D = 41.66$*,* $F_R = 30.16$*) using random LD-clumping. Using combined data, an average of 55 instruments are used (*$F_C = 55.17$*) using standard LD-clumping whereas an average of 175 instruments are used (*$F_C = 41.09$*) using random LD-clumping.* . . . . . . . . . . . . . . . . 8

Supplementary tables S2, S3, S4, S5, S6, S7, and S9 are too large to be included in this document and are available in an `xlsx` file.

### Supplementary Figures

S1 *Effect of the total exposure sample size used for discovery on naive IVW (light blue), IVW using a 3-sample design (dark blue), MR-RAPS using a 3-sample design (dark purple), MR-RAPS using ZP (light purple), MR-RAPS using UMVCUE (salmon pink) and IVW using the regression-calibration approach (brick red) in presence of balanced pleiotropy (mean effect, panel A) and RMSE, panel B), across 1,000 simulations). Instruments are selected using a p-value threshold of* $5\times10^{-8}$ *(*$T \approx 5.45$*) and the true causal effect (0.2) is indicated by the dashed line in panel A).* . . . . . . . . . . . . . . . . . . . . . . . . . . . . . . . . . . 9

S2 *Effect of the outcome sample size used on naive IVW (light blue), IVW using a 3-sample design (dark blue), MR-RAPS using a 3-sample design (dark purple), MR-RAPS using ZP (light purple), MR-RAPS using UMVCUE (salmon pink) and IVW using the regression-calibration approach (brick red) in presence of balanced pleiotropy (mean effect, panel A) and RMSE, panel B), across 1,000 simulations). Instruments are selected using a p-value threshold of* $5 \times 10^{-8}$ *(*$T \approx 5.45$*) and the true causal effect (0.2) is indicated by the dashed line in panel A).* . . 9

S3 *Effect of the selection threshold on naive IVW (light blue), IVW using a 3-sample design (dark blue), MR-RAPS using a 3-sample design (dark purple), MR-RAPS using ZP (light purple), MR-RAPS using UMVCUE (salmon pink), IVW using the regression-calibration approach (brick red), simple median-based using the regression-calibration approach (dark orange, dashed), weighted median-based using the regression-calibration approach (dark orange, dotted), simple mode-based using the regression-calibration approach (light orange, dashed), weighted mode-based using the regression-calibration approach (light orange, dotted) in presence of weak (panel A) and strong (panel B) correlated pleiotropy (mean effect across 1,000 simulations). Instruments were selected with different selection thresholds (T) and the true causal effect (0.2) is indicated by the dashed line.* . . . . . . . . . . . . . . . . . . . . . . . . 10

S4 *Comparison of results using standard LD-clumping and random LD-clumping on naive IVW (light blue), MR-RAPS using a 3-sample design (dark purple), and MR-RAPS using ZP (light purple) (from 100 random samplings). Instruments are selected using a p-value threshold of* $5 \times 10^{-8}$ *(*$T \approx 5.45$*) and the true causal effect is expected to be 1 (dashed line)* . . . . . . . . . . . . . . . . . . . . . . . . . . . . . 11

S5 *LocusZoom Plot for one region on chromosome 3, illustrating the instrument selection process for BMI (for one of the random samplings) in presence of LD. Several genetic instruments are passing the genome-wide significance threshold (grey dashed line) but they are all highly correlated (LD* $r^2 > 0.8$*) with the lead SNP in the region (rs1007141, in purple). Using a standard LD-clumping approach only this SNP will be kept as an instrument for MR, but to be used, it actually had to pass a threshold of* $4.70 \times 10^{-9}$ *(second most significant association in the region) and not* $5 \times 10^{-8}$. . . . . . . . . . . . . . . . . . . . . . . . . . 12

| Approach | SNP selection | SNP-exposure association | Robust to… | Remarks |
|---|---|---|---|---|
| **Naive IVW** | Discovery + Replication | Discovery + Replication | -<br>- | |
| **IVW, 3 sample design** | Discovery | Replication | Winner's curse<br>- | |
| **MR-RAPS, 3 sample design** | Discovery | Replication | Winner's curse, weak instruments | |
| **MR-RAPS, ZP** | Discovery + Replication | ZP ( Discovery+ Replication ) | Winner's curse, weak instruments | |
| **MR-RAPS, UMVCUE** | Discovery | UMVCUE ( Discovery + Replication ) | Winner's curse, weak instruments | |
| **Regression calibration** | Discovery | Replication | Winner's curse, weak instruments | possibility to use pleiotropy-robust methods |
| **MRlap** | Discovery + Replication | Discovery + Replication | Winner's curse, weak instruments | possibility to use overlapping samples |

Table S1: *Summary of the data used by each approach for SNP selection and SNP-exposure association estimation.*

| Estimator | Mean absolute bias<br>standard LD-clumping | <br>random LD-clumping |
|---|---|---|
| Naive IVW | 0.160 | 0.180 |
| MR-RAPS, ZP | 0.059 | 0.029 |
| MR-RAPS, UMVCUE | 0.020 | 0.010 |

Table S8: *Mean absolute bias for naive IVW, MR-RAPS using ZP and MR-RAPS using UMVCUE when estimating the causal effect of BMI on itself, comparing two LD-clumping approaches. Instruments were selected using a p-value threshold of* $5 \times 10^{-8}$ *(*$T \approx 5.45$*). Using discovery data, an average of 55 instruments are used (*$F_D = 51.26$*,* $F_R = 37.64$*) using standard LD-clumping whereas an average of 55 instruments are used (*$F_D = 41.66$*,* $F_R = 30.16$*) using random LD-clumping. Using combined data, an average of 55 instruments are used (*$F_C = 55.17$*) using standard LD-clumping whereas an average of 175 instruments are used (*$F_C = 41.09$*) using random LD-clumping.*

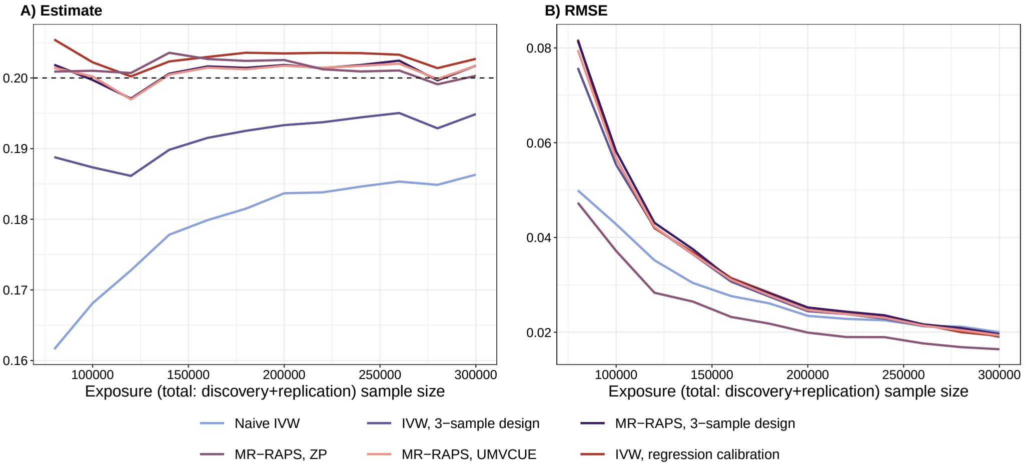


Figure S1: *Effect of the total exposure sample size used for discovery on naive IVW (light blue), IVW using a 3-sample design (dark blue), MR-RAPS using a 3-sample design (dark purple), MR-RAPS using ZP (light purple), MR-RAPS using UMVCUE (salmon pink) and IVW using the regression-calibration approach (brick red) in presence of balanced pleiotropy (mean effect, panel A) and RMSE, panel B), across 1,000 simulations). Instruments are selected using a p-value threshold of* $5 \times 10^{-8}$ *(*$T \approx 5.45$*) and the true causal effect (0.2) is indicated by the dashed line in panel A).*

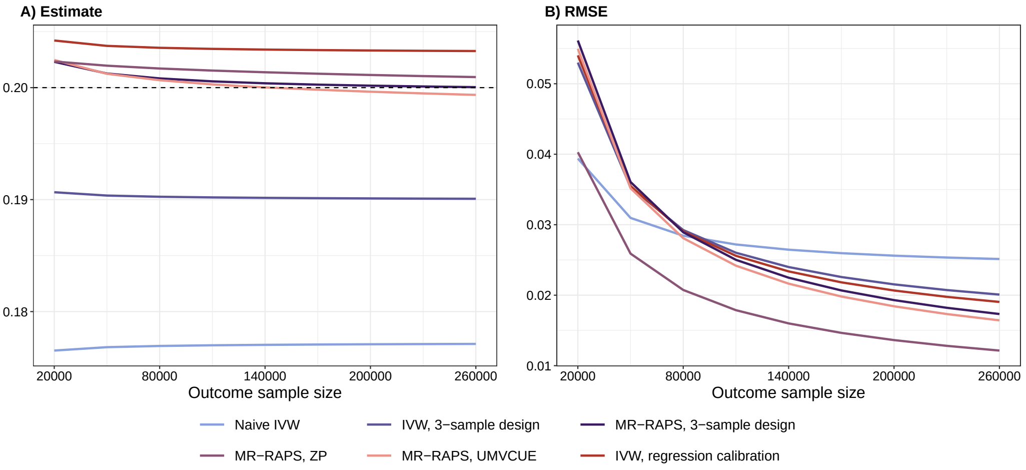


Figure S2: *Effect of the outcome sample size used on naive IVW (light blue), IVW using a 3-sample design (dark blue), MR-RAPS using a 3-sample design (dark purple), MR-RAPS using ZP (light purple), MR-RAPS using UMVCUE (salmon pink) and IVW using the regression-calibration approach (brick red) in presence of balanced pleiotropy (mean effect, panel A) and RMSE, panel B), across 1,000 simulations). Instruments are selected using a p-value threshold of* $5 \times 10^{-8}$ *(*$T \approx 5.45$*) and the true causal effect (0.2) is indicated by the dashed line in panel A).*

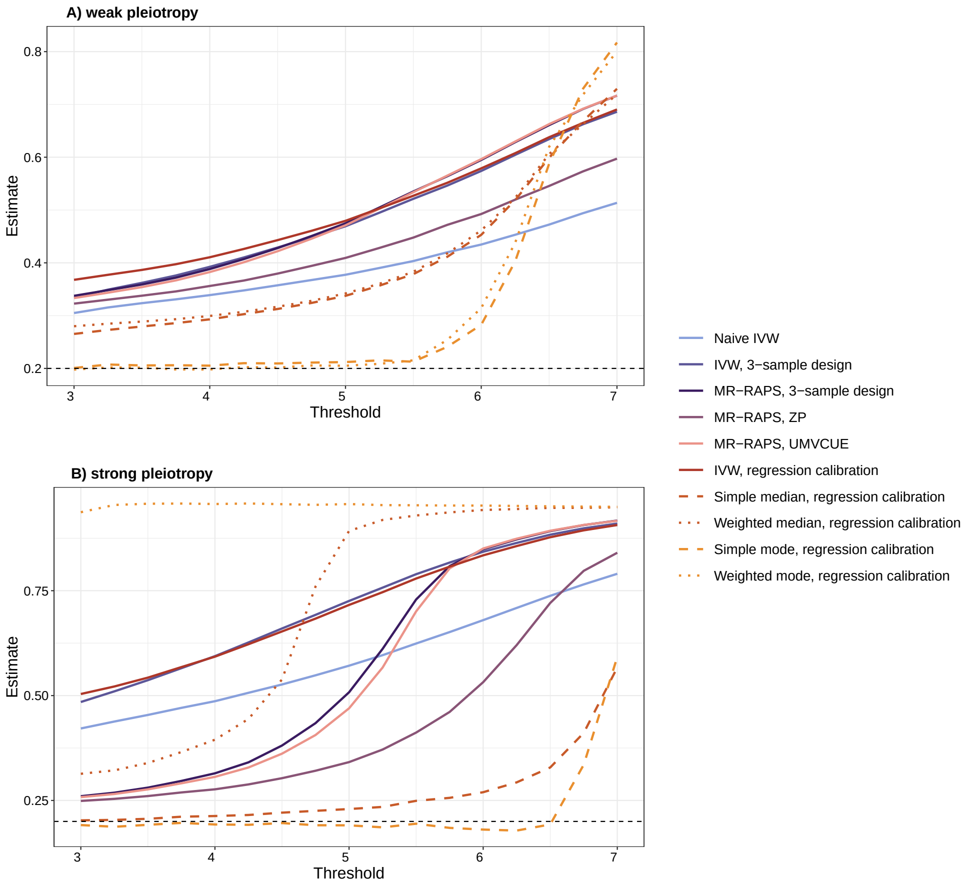


Figure S3: *Effect of the selection threshold on naive IVW (light blue), IVW using a 3-sample design (dark blue), MR-RAPS using a 3-sample design (dark purple), MR-RAPS using ZP (light purple), MR-RAPS using UMVCUE (salmon pink), IVW using the regression-calibration approach (brick red), simple median-based using the regression-calibration approach (dark orange, dashed), weighted median-based using the regression-calibration approach (dark orange, dotted), simple mode-based using the regression-calibration approach (light orange, dashed), weighted mode-based using the regression-calibration approach (light orange, dotted) in presence of weak (panel A) and strong (panel B) correlated pleiotropy (mean effect across 1,000 simulations). Instruments were selected with different selection thresholds ($T$) and the true causal effect (0.2) is indicated by the dashed line.*

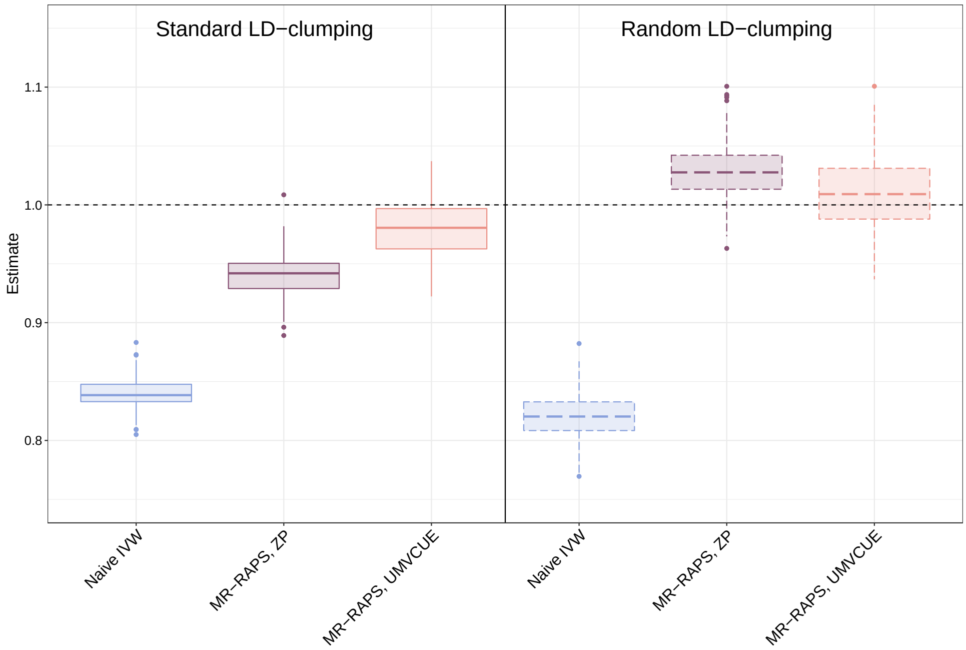


Figure S4: *Comparison of results using standard LD-clumping and random LD-clumping on naive IVW (light blue), MR-RAPS using a 3-sample design (dark purple), and MR-RAPS using ZP (light purple) (from 100 random samplings). Instruments are selected using a p-value threshold of* $5 \times 10^{-8}$ *(*$T \approx 5.45$*) and the true causal effect is expected to be 1 (dashed line)*

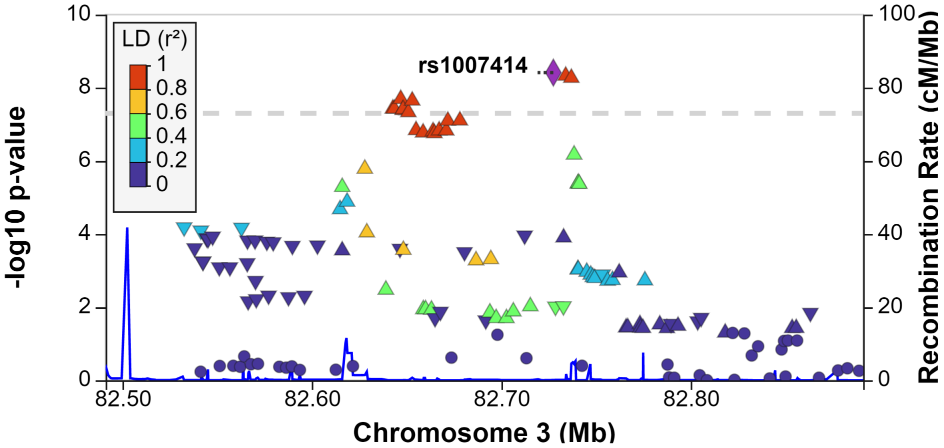


Figure S5: *LocusZoom Plot for one region on chromosome 3, illustrating the instrument selection process for BMI (for one of the random samplings) in presence of LD. Several genetic instruments are passing the genome-wide significance threshold (grey dashed line) but they are all highly correlated (LD* $r^2 > 0.8$*) with the lead SNP in the region (rs1007141, in purple). Using a standard LD-clumping approach only this SNP will be kept as an instrument for MR, but to be used, it actually had to pass a threshold of* $4.70 \times 10^{-9}$ *(second most significant association in the region) and not* $5 \times 10^{-8}$.
